# Differential Glutamatergic Inputs to Semilunar Granule Cells and Granule Cells Underscore Dentate Gyrus Projection Neuron Diversity

**DOI:** 10.1101/2025.03.14.643192

**Authors:** Laura Dovek, Chia-Wei Yeh, Anh-Tho Nguyen, Emmanuel Green, Vijayalakshmi Santhakumar

**Author notes:** Correspondence: Vijayalakshmi Santhakumar, PhD Department of Molecular, Cell and Systems Biology University of California, Riverside 3401 Watkins Drive (Rm 2111 Biological Sciences Building) Riverside, CA 92521. Department of Neurology, Oregon Health & Science University (Laura Dovek).

## Abstract

Semilunar Granule Cells (SGCs) are sparse dentate gyrus projection neurons whose role in the dentate circuit, including pathway-specific inputs, remains unknown. We report that SGCs receive more frequent spontaneous excitatory synaptic inputs than granule cells (GCs). Dual GC-SGC recordings identified that SGCs receive stronger medial entorhinal cortex and associational synaptic drive but lack short-term facilitation of lateral entorhinal cortex inputs observed in GCs. Consistently, the probability of firing in response to medial perforant path input was greater in SGCs than in GCs. Additionally, the dendritic spine density on proximal and middle dendrites was higher in SGCs than GCs. However, the strength of commissural and supramammillary nuclear inputs and dendritic input integration, examined in passive morphometric simulations, did not differ between cell types. Activity-dependent labeling identified an overrepresentation of SGCs among neuronal ensembles in both mice trained in a spatial memory task and task-naïve controls. The divergence of modality-specific entorhinal inputs to SGCs and GCs identified here can enable parallel processing of information streams and expand the computational capacity of the dentate gyrus.

## Introduction

The hippocampus plays a critical role in forming and consolidating new memories. In particular, the dentate gyrus (DG) is proposed to regulate the flow and integration of multisensory input streams to the hippocampus and promote the formation of distinct memory traces. Studies in the hippocampus and cortex have revealed that seemingly homogeneous neurons can exhibit subtle diversity in structure, intrinsic physiology, and connectivity, which can enhance the information processing in the circuit (Slomianka et al., 2011; Soltesz and Losonczy, 2018). Recent studies have identified a subtype of DG projection neurons, semilunar granule cells (SGCs), with several features that distinguish them from the classical granule cells (GCs). SGCs were first identified by their location above the granule cell layer (GCL), their wide dendritic spread, and characteristic “semilunar” somatic shape (Williams et al., 2007; Gupta et al., 2012) and defined as a distinct subpopulation on the basis of unbiased morphometric analysis (Gupta et al., 2020; Afrasiabi et al., 2022). Functionally, compared to GCs, SGCs have lower input resistance and less adapting firing pattern (Williams et al., 2007; Afrasiabi et al., 2022; Dovek et al., 2025). SGCs receive more frequent inhibitory postsynaptic currents (IPSCs) and have higher extrasynaptic GABA currents (Gupta et al., 2012; Gupta et al., 2022) indicating differences in circuit connectivity. Similar to CA1 pyramidal cell subtypes, which are generated in different waves of neurogenesis (Bayer, 1980), SGCs are preferentially generated during late embryonic generation of GC development (Save et al., 2019). Although SGCs are a relatively sparse subtype (Save et al., 2019) they are disproportionately labeled as part of activity-dependent behavioral engrams, suggesting a preferential role in dentate processing (Erwin et al., 2020; Dovek et al., 2025; Dovek et al., 2026). However, it remains unclear whether SGCs and GCs differ in their glutamatergic synaptic inputs.

Afferent drive to the DG segregates into layered and functionally distinct input streams that are integrated in GC and SGC dendrites to evoke firing. Massive cortical inputs from the entorhinal cortex (EC) impinge on the DG through the perforant path (PP) afferent fibers. The PP segregates into two stratified input streams with distinct information content: the medial perforant path (MPP) arising from the medial entorhinal cortex (MEC) projects to GC and SGC dendrites in the middle molecular layer (MML), and the lateral perforant path (LPP) arising from the lateral entorhinal cortex (LEC) targets distal dendrites in the outer molecular layer (OML). While MEC is involved in processing spatial (where/context) information, the LEC carries information about the object (what/content) (Knierim et al., 2006; Knierim et al., 2014; Fernández-Ruiz et al., 2021). While prior studies in rats indicate that PP-driven excitatory currents do not differ between GCs and SGCs (Williams et al., 2007), it remains unclear whether mass electrical activation obscures pathway-specific differences. Apart from PP inputs, axons of glutamatergic mossy cells (MCs) in the DG hilus extend robust commissural and associational projections along the septotemporal axis of the DG, terminating on the contralateral and ipsilateral GC dendrites, respectively (Buckmaster et al., 1996; Houser et al., 2020; Botterill et al., 2021). Ventral mossy cells project dense inputs to the proximal dendrites of GCs in the inner molecular layer (IML). In ventral hippocampal slices, stimulation of associational inputs in the hilus elicits larger glutamatergic synaptic currents in SGCs than in GCs (Williams et al., 2007). However, recent studies have identified that dorsal mossy cell terminals are more diffusely distributed and overlap with MEC inputs in the MML (Houser et al., 2020; Botterill et al., 2021). Ventral mossy cells respond to environmental novelty, such as context discrimination, and preferentially project to dorsal GCs (Fredes et al., 2021). Yet, little is known about the functional characteristics of commissural dorsal and ventral mossy cell projections to dentate cell types. In addition, the supramammillary nucleus (SuM) of the hypothalamus, implicated in spatial learning, projects to the supragranular layer of the DG (Hashimotodani et al., 2018; Ajibola et al., 2021), where it could innervate GCs and SGCs. It is currently unknown whether these input streams differentially influence SGCs and GCs.

Neuronal activation is determined both by the strength of the dendritic inputs and by how the inputs are integrated in the dendrites. Integration and attenuation of inputs are shaped by cell morphology and dendritic active and passive membrane properties. In this regard, GCs have been found to demonstrate unique dendritic input propagation with limited input attenuation in the distal and mid dendritic segments and robust signal attenuation at the proximal dendrite, not observed in other excitatory neurons (Krueppel et al., 2011). Computational studies have identified that morphological features of GCs contribute to their unique dendritic integration and attenuation characteristics. Given the morphological differences between GCs and SGCs, it is important to determine whether neuronal structure affects input handling in the two cell types.

We examined the spontaneous glutamatergic drive and pathway-specific inputs to GCs and SGCs. We find that SGCs receive significantly more spontaneous excitatory synaptic inputs than GCs. Optically driven recordings of excitatory synaptic inputs to GC-SGC pairs identified cell-type and pathway-specific differences in strength and short-term plasticity of entorhinal cortex inputs to SGCs and GCs. Simulations in morphologically constrained detailed multicompartmental models of GCs and SGCs showed that dendritic input integration did not differ between the cell types. Activity-dependent labeling of neurons revealed greater proportional recruitment of SGCs to neuronal ensembles both in task-naïve control mice in their home cage and in littermate mice trained in a dentate-dependent spatial memory task. Our results extend the known differences between GCs and SGCs to their circuit connectivity and support the proposal that SGCs and GCs form parallel interacting microcircuits, and are positioned to enhance information processing in the DG.

## Materials and Methods

### Animals

All experiments were conducted under IACUC protocols approved by the University of California at Riverside and conformed with ARRIVE guidelines. Male and female C57Bl6/J mice (p28-35) were used in slice electrophysiology studies. Inducible Fos-cre mice (TRAP2: Fostm2.1^(icre/ERT2)Luo/J^; Jackson Laboratories #030323) were back-crossed with C57BL6/N and were bred with reporter line tdT-Ai14 mice (B6;129S6-Gt(ROSA)^26Sortm14(CAG-tdTomato)Hze/J^; Jackson Laboratories # 007908) to create TRAP2-tdT mice for behavior experiments. Male and female TRAP2-tdT mice four to eight weeks old were used in behavior experiments. Mice were housed with littermates (up to 5 mice per cage) in a 12/12 h light/dark cycle. Food and water were provided ad libitum.

### Stereotaxic Injections

Male and female C57Bl6/J mice were injected with AAV in the lateral and medial entorhinal cortex, hilus (dorsal and ventral), and the supramammillary nucleus (SuM) to selectively label specific glutamatergic inputs to GCs and SGCs. Juvenile mice (p28-35) were anesthetized with isoflurane and secured in a small animal stereotaxic frame (Kopf). The scalp was injected (s.c.) with bupivacaine (2 mg/kg) at the site of incision. Animals also received a subcutaneous dose of carprofen (5 mg/kg) and buprenorphine (Ethiqa XR at 3.25 mg/kg) at the start of surgery for analgesia. A 10 µl Hamilton syringe was used with a Harvard Apparatus (11 Elite Nanomite) syringe pump for injections.

Stereotaxic coordinates (in mm) used for the entorhinal cortex regions: anterior/posterior (AP) in relation to true lambda; medial/lateral (ML) in relation to midline; and dorsal/ventral (DV) in relation to the dorsal brain surface. MEC: AP: -0.7, ML: 3.4, DV: -2.5. LEC: AP: + 0.8, ML: 4.3, DV: -3.6 (Donohue et al., 2021). To label MPP and LPP projections, mice were bilaterally injected with 180 nl at 25 nl/min of AAV-hSyn-hChR2(H134R)-EYFP (AAV5) or AAV-hSyn-EGFP (AAV5) for controls at a titer of 1-5 x10^13^ vg/ml.

Mice in which the dorsal or ventral hilus were targeted received unilateral injections of AAV-CaMKIIa-hChR2(H134R)-EYFP (AAV5) or AAV-CaMKIIa-EGFP (AAV5) for controls (300 nl at 25 nl/min, at a titer of 10^13^ vg/ml) diluted in 1:1 sterile saline. The following hilar coordinates, AP is in relation to bregma were adopted: Dorsal hilus, AP: -2.1, ML: -1.25, DV: - 1.9 from skull surface. Ventral hilus, AP: -3.4, ML: -2.7, DV: -3.4 from skull surface (adapted from Botterill et al., 2021). For SuM injections, mice received bilateral injections of AAV5-hSyn-hChR2(H134R)-EYFP (180 nl at 25 nl/min) at the following coordinates: AP -2.9 mm, ML ±0.15 mm, DV -4.9 mm from skull surface (Ajibola et al., 2021).

Injection site and projections with GFP and eYFP were visualized under epifluorescence illumination (λ=505 nm) using a Thorlabs 4-wavelength high-power LED source. Dual recordings from SGC-GC pairs were adopted to limit confounding effects of variable AAV labeling and obtain optically evoked EPSCs. pAAV-hSyn-hChR2(H134R)-EYFP and pAAV-CaMKIIa-hChR2(H134R)-EYFP were developed by Karl Deisseroth (Addgene viral prep # 26973-AAV5 & # 26969-AAV5; http://n2t.net/addgene:26973 & http://n2t.net/addgene:26969; RRID:Addgene_26973 & RRID:Addgene_26969, Lee et al., 2010; Chan et al., 2017) pAAV-hSyn-EGFP and pAAV-CaMKIIa-EGFP were developed by Bryan Roth (Addgene viral prep # 50465-AAV5 & Addgene viral prep # 50469-AAV5; http://n2t.net/addgene:50465 & http://n2t.net/addgene:50469; RRID:Addgene_50465 & RRID:Addgene_50469)

### Slice Physiology

Naïve C57Bl6/J mice (p28-35), or C57Bl6/J mice 3-4 weeks following AAV injection were euthanized under isoflurane anesthesia for preparation of horizontal brain slices (300 μm) using a Leica VT1200S Vibratome in ice cold sucrose artificial cerebrospinal fluid (sucrose-aCSF) containing (in mM): 85 NaCl, 75 sucrose, 24 NaHCO_3_, 25 glucose, 4 MgCl_2_, 2.5 KCl, 1.25 NaH_2_PO_4_, and 0.5 CaCl_2_. Slices from the ventral-mid hippocampus were bisected and incubated at 32°C for 30 min in a holding chamber containing an equal volume of sucrose-aCSF and recording aCSF, and were subsequently held at room temperature for an additional 30 min before use. The recording aCSF contained (in mM): 126 NaCl, 2.5 KCl, 2 CaCl_2_, 2 MgCl_2_, 1.25 NaH_2_PO_4_, 26 NaHCO_3_, and 10 glucose. All solutions were saturated with 95% O_2_ and 5% CO_2_ and maintained at a pH of 7.4 for 2-6 hours (Gupta et al., 2012; Yu et al., 2015; Afrasiabi et al., 2022). Slices were transferred to a submerged recording chamber and perfused with oxygenated aCSF at 33°C. Whole-cell voltage-clamp and current-clamp recordings from GCs in the GCL and presumed SGCs in the IML or edge of the GCL were performed under IR-DIC visualization with Nikon Eclipse FN-1 (Nikon Corporation) using 40X water-immersion objective. Recordings were obtained using filamented borosilicate microelectrodes (3-7 MΩ) and recorded using Molecular Devices MultiClamp 700B amplifier. Data were low-pass filtered at 2 kHz, digitized using Axon DigiData 1400A (Molecular Devices), and acquired using pClamp11 at 10 kHz sampling frequency. Recordings were performed using K-gluconate internal solution (K-gluc) containing (in mM) 126 K-gluconate, 4 KCl, 10 HEPES, 4 Mg-ATP, 0.3 Na-GTP, and 10 phosphocreatine or cesium methane sulfonate (CsMeSO_4_) internal solution (cesium) containing 140 cesium methane sulfonate, 10 HEPES, 5 NaCl, 0.2 EGTA, 2 Mg-ATP, and 0.2 Na-GTP. (pH 7.25; 270–290 mOsm). Biocytin (0.2 %) was included in the internal solution for *post hoc* cell identification (Gupta et al., 2012; Yu et al., 2015; Afrasiabi et al., 2022).

Spontaneous and electrically evoked EPSCs were recorded using the cesium-based internal solution with cells held at -70 mV in voltage clamp to isolate EPSCs. Evoked EPSCs were elicited by single stimulation of either the perforant path or the hilus using FHC stimulating electrodes placed at the junction of the dorsal blade and the crest just outside the fissure for PP (Korgaonkar et al., 2020) or in the hilus, avoiding the GCL and CA3 pyramidal cells, for hilar stimulation. An Iso-Flex (AMPI Israel) stimulating box was used to deliver stimuli at 0.2 mA increments (3 sweeps, 50 µs pulse duration).

For experiments involving AAV-driven optically evoked PSCs, ChR2-expressing axonal terminals were optically stimulated using wide-field blue (λ=470 nm, 0.9 mW) illumination of the full field under a 40X objective using a Thorlabs 4-wavelength High-Power LED source at a single saturating stimulus intensity. Cells received a 10 ms light pulse for 10 sweeps at 500 ms intervals or a 10 Hz train of light pulses with a 10 ms pulse width. Cell-attached recordings (Wei et al., 2021) and whole-cell voltage-clamp (Afrasiabi et al., 2022; Dovek et al., 2025) were performed on dentate GCs and SGCs to investigate the recruitment and synaptic strength of either MEC or LEC inputs using K-gluconate-based internal solution. Whole-cell recordings were discontinued if series resistance increased by >20% or if access resistance was over 25 MΩ. Post hoc biocytin immunostaining and morphological analysis were used to definitively identify all SGCs and GCs included in this study.

### Immunostaining, Cell Morphology and Reconstructions

Following physiological recordings, slices were fixed in 0.1mM phosphate buffer containing 4% paraformaldehyde at 4°C overnight. Slices were washed with PBS and then incubated in Alexa Fluor® 594-conjugated streptavidin (1:1000 Thermo Fisher, S11227) in PBS with 0.3% Triton X-100 for 2 hours at room temperature.

After behavioral experiments, animals underwent transcardial perfusion with PBS followed by a 4% paraformaldehyde solution, under Euthasol anesthesia. Brains were extracted and stored in 4% paraformaldehyde at a temperature of 4°C for 3 hours before being transferred to PBS. The brains were sliced coronally into 50 µm sections using a VF-310-0Z Compresstome (Precisionary Instruments); and 5 sequential sections were chosen, each 250 µm apart across the septotemporal axis. Free-floating sections were blocked in 10% goat serum in PBS with 0.3% Triton X-100 for one hour. Sections were incubated in 4°C overnight in primary antibody for c-Fos (1:750, Rabbit mAb Cell Signaling Technology, cat #2250). The following day, sections were incubated in goat anti-rabbit Alexa Fluor® 488 secondary antibody (1:500 Abcam, cat #150077) for 1 hour and were then mounted on glass slides using Vectashield®.+DAPI mounting media (Dovek et al., 2025).

Sections were imaged using a Zeiss Axioscope-5 with Stereo Investigator (MBF Bioscience) for analysis. Cell counts and cell type classification and evaluation of double labeling were conducted by experimenters blinded to treatments. Cells with compact dendritic arbors and somata with greater length than width were classified as GCs, and those with wide dendritic angle and 2 or more primary dendrites and greater somatic width than height were classified as SGCs (Gupta et al., 2020; Afrasiabi et al., 2022; Dovek et al., 2025).

Following *post hoc* immunostaining for biocytin recovery, a subset of biocytin-filled cells was imaged using a Zeiss 800 Upright Microscope with Airyscan Fast using 20X lens at 0.23 μm steps. Dendritic segments were imaged at higher resolution using a 40X oil lens at 0.04 μm steps. Images were obtained at three regions along the cell: closest to the soma, the middle and the end of the dendrites for proximal, middle, and distal dendrites, respectively. Spines within each region were selected at random based on optimal depth for imaging and reconstructed from 3D image stacks using Neurolucida 360. Reconstructions were user-guided and began with initial tree construction in which all trees with complete imaging from all directions (checked through 3D rotation of the image) were reconstructed. Subsequently, using the user-guided spine detection tool, each spine was selected by the user and reconstructed by the software. Automated spine classifications were used, followed by experimenter verification. Counts and classifications were extracted using Neurolucida explorer. Spines were classified into canonical types (stubby, mushroom, thin, filopodial) in Neurolucida 360 based on the method described previously (Rodriguez et al., 2008). Spine types were determined using a combination of measured values (spine length, head and neck diameter) and derived ratios (Aspect ratio defined as the ratio of the depth from tip to widest point of spine and the width of spine at widest point). Due to variability in the length of dendritic segments imaged, spine densities were calculated by dividing the total number of spines from a segment by the total length of that segment.

### Simulations

Granule cells and semilunar granule cells imaged using the Zeiss 800 Upright microscope were imported and reconstructed on Neurolucida 360. Following reconstructions, ASCII files were exported into “Trees” software (Cuntz et al., 2011) to enable segmenting the dendrites into functionally distinct thirds representing the IML, MML, and OML. Segmenting was performed following resampling to make the distance between all nodes equal. Using “PVEC, PathLength” function, the dendrites were split into three segments and assigned titles of IML, MML, and OML accordingly. Segmented neuronal morphologies were imported into NEURON (Hines and Carnevale, 2001) to insert conductance mechanisms. Active and passive conductances were distributed as described previously with a minor modification (Table 1). GC and SGC somata were simulated as cylindrical compartments (in µm, GC: Length 16.8, Dia: 16.8; SGC: Length: 16.8, Dia 18.3). Dendritic channel densities were based on previously described parameters (Santhakumar et al., 2005) and maintained identical in all dendritic compartments (Table 1).

**TABLE 1.**
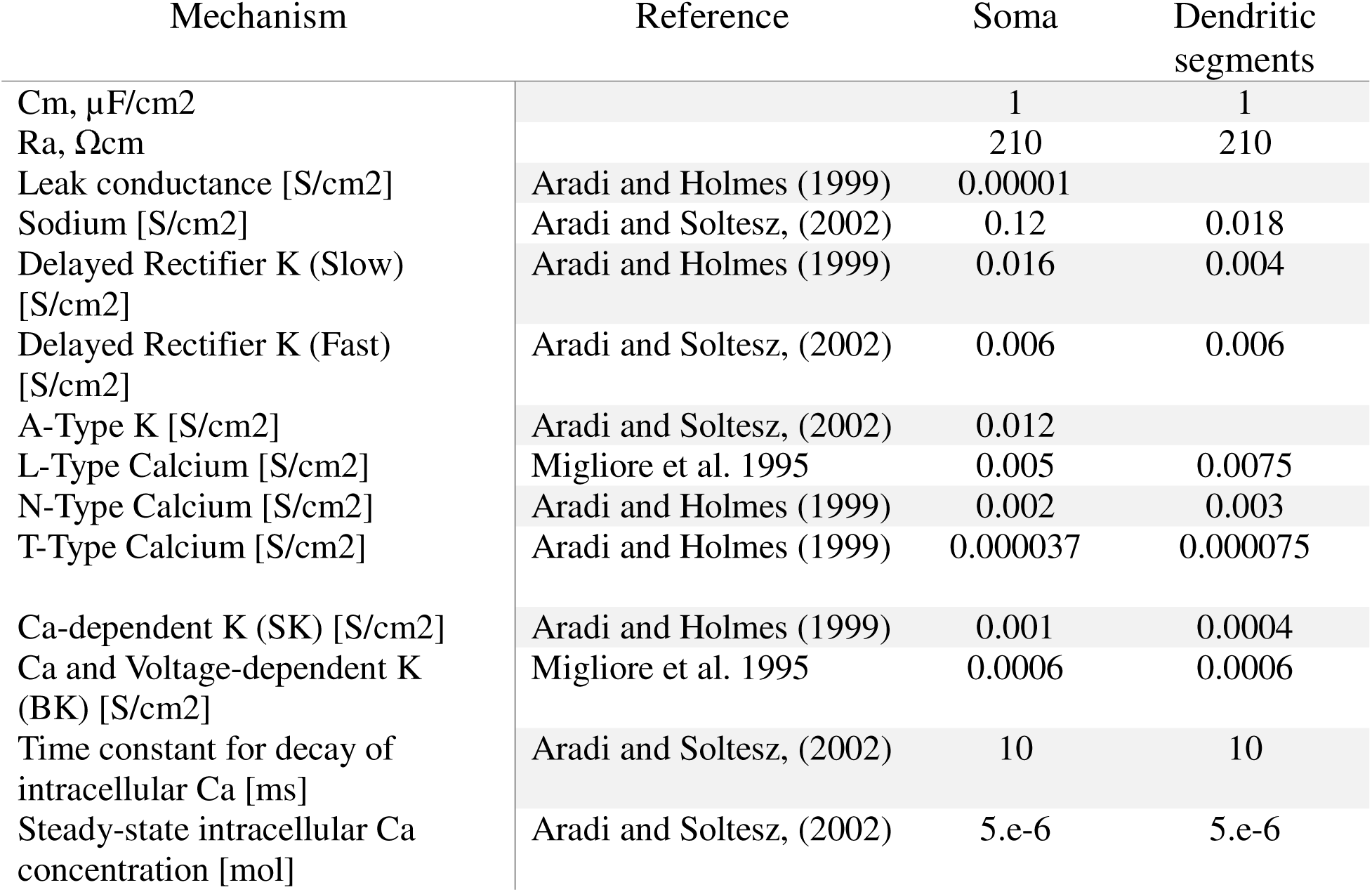
Passive parameters and maximum conductance of the channels in model cells.

For experiments examining EPSP amplitude attenuation, a single, identical AMPA synaptic conductance (Golding et al., 2005) was located in the appropriate dendritic segment (IML, MML, or OML) and activated with an artificial spike. The resulting EPSP was measured both at the local dendrite and at the soma. Attenuation was evaluated by computing the EPSP peak amplitude at the soma as a percent of the local dendrite peak EPSP amplitude. A total of 9 trials with synapses located pseudo-randomly in different compartments within each of the three dendritic segments (IML, MML, and OML) were simulated in each reconstructed neuron.

In experiments evaluating input summation, 5 AMPA synaptic conductances were randomly placed within the MML segments of the cell’s dendrites. Five independent simulations were conducted with synaptic conductance distributed either on the same branch or with the 5 synaptic conductances randomly distributed across branches in the middle dendritic segment. A total of 10 trials per reconstruction were implemented, and the peak EPSP amplitude at the soma was compared between cell types.

### 4-Hydroxy Tamoxifen Preparation

4-Hydroxy Tamoxifen was dissolved in 100% ethanol at a concentration of 20 mg/mL by sonicating the solution at 37°C for 30 minutes or until it was fully dissolved. The solution was then aliquoted and stored at -20°C. On the day of use, 4-OHT was redissolved by sonicating the solution at 37°C for 10 minutes. A 1:4 mixture of castor oil and sunflower seed oil was added respectively to give a final concentration of 10 mg/mL. The remaining ethanol in solution was evaporated by speed vacuuming in a centrifuge. Animals received 4-OHT (50 mg/kg) intraperitoneally (DeNardo et al., 2019; Dovek et al., 2025).

### Barnes Maze

Male and female TRAP2-tdT mice were trained in a Barnes maze spatial learning task. A cohort of littermate controls went through all handling and activities other than being exposed to maze training. The Barnes maze table is 92 cm in diameter and has 20 holes that are 5 cm in diameter. A false floor with a removable escape box could be swapped for an additional false floor piece. The maze was set up in the middle of 4 curtain walls with two bright lights and a camera above the maze for recording. Each curtain had a different set of visual cues (various shapes cut from felt). The escape hole was positioned between two visual cues. Animals were kept outside of the curtain in a dark room until their turn to run the trial.

Three days before testing, animals were handled for about a minute daily to habituate animals to the experimenter. One day (Day 0) before beginning behavior, animals were individually housed because this mouse line is hyperactive and to prevent confounding results. Animals were brought to the behavior room and left alone to habituate in their cages for at least 1 hour before beginning training. Additional habituation was performed on day 1 of training, during which animals were placed in the starter cup on the table for 1 minute. They were then guided towards an escape box in a temporary location different from the experimental location.

During the initial 1-6 days of acquisition training, each animal in the experimental group performed three 180-second trials. For animals in the active experimental group, the experimenter performed each trial for all animals before starting the next trial with a minimum inter-trial interval of 15 minutes. Mice that did not enter the escape box within 180 seconds were guided by the experimenter to the escape box and returned to their home cage. On day 6 of training, the animals were brought into the room 5 hours before testing. 10 minutes before Barnes maze testing, each mouse received 4-hydroxy tamoxifen (50 mg/kg, i.p). Animals were left in the room for an additional 5 hours after testing to limit neuronal activity labeling not related to the behavioral paradigm. During the probe trial on day 7, the escape box was replaced to match the other holes. The table was rotated 180 degrees to account for potential olfactory cues. Each animal explored the table for 90 seconds before being returned to the home cage. A week following induction (6 days following the probe trial) on experimental day 12-13, animals repeated the training to reactivate the memory engram. Animals were sacrificed 90 minutes after performing the task on day 13 for immunostaining.

Between each trial, the entire table and escape box were wiped down with 70% ethanol and allowed to dry completely. Behavioral data were analyzed using Any-Maze software by a blinded experimenter and using the automated BUNS analysis software (Illouz et al., 2016).

### Analysis

Analysis of optical and stimulus-evoked EPSCs as well as spontaneous EPSCs (sEPSCs) was conducted using EasyElectrophysiology software (Easy Electrophysiology Ltd). The evoked PSC peak was measured from the baseline to the first peak occurring within 4 ms of the start of the stimulus. sEPSCs were detected using a threshold search algorithm following a baseline period of 180 seconds and events were confirmed by the experimenter. Any “noise” that spuriously met trigger specifications was rejected. Instantaneous frequency was calculated from the first 100 accepted events. The reciprocal of inter-event interval (IEI) values generated in EasyElectrophysiology were used to compute instantaneous event frequencies used in further analysis. Amplitude, IEI Charge transfer (Area Under the curve, pA • ms), Rise Time (ms), and Biexponential Fit of Decay (ms) from the first 100 accepted events from each cell were used to generate a cumulative probability distribution from comparison across cell types. A bi-exponential fit was used when calculating decay. Rise time was calculated between 10 and 90% of the event amplitude.

### Quantification and Statistical Analysis

Sample sizes were not predetermined and conformed with those employed in the field. Significance was set to p<0.05, subject to appropriate Bonferroni Correction. Statistical analysis and figure generation were performed using GraphPad Prism 10. Data were tested for normality using the Shapiro-Wilk test, and data that failed normality were analyzed using non-parametric tests. Non-parametric unpaired Kolmogorov-Smirnov (KS) test, unpaired Mann-Whitney test, Wilcoxon matched-pairs signed-rank test, and parametric nested t-test, two-way ANOVA with Holm-Sidak post hoc comparison, repeated measures ANOVA, Kruskal-Wallis ANOVA with Dunn’s multiple comparisons were used as appropriate. Data are presented as mean ± SEM. Additional graphics were created using Biorender.

## Results

### Semilunar granule cells receive more frequent spontaneous glutamatergic inputs

Recent studies have identified differences in intrinsic physiology and inhibitory synaptic inputs between GCs and SGCs (Williams et al., 2007; Gupta et al., 2012; Gupta et al., 2020; Afrasiabi et al., 2022; Dovek et al., 2025). However, little is known about the steady-state excitatory drive to SGCs. To address this knowledge gap, we recorded spontaneous excitatory postsynaptic currents (sEPSCs) in GCs and SGCs from the ventral hippocampus in horizontal brain slices. To target mature GCs, recordings were obtained from the middle and outer thirds of the granular cell layer as in prior studies (Gupta et al., 2020; Afrasiabi et al., 2022; Dovek et al., 2025; Dovek et al., 2026). SGCs were identified by the presence of multiple primary dendrites, a greater dendritic angle, as well as soma location and a larger soma aspect ratio compared to GCs, as described previously (Gupta et al., 2020; Afrasiabi et al., 2022) (Fig. 1A-C). A summary plot of the number of primary dendrites and dendritic angle in the recorded neurons revealed two groups, aligning with experimenter-defined cell categories, GCs (black) and SGCs (blue) (Fig. 1C), confirming the distinct structures of GCs and SGCs. Any cell not clearly categorized into either group was excluded from further analysis. Current recordings from a holding potential of - 70 mV showed that SGCs received more frequent sEPSCs than GCs (Fig. 1D,E, frequency in Hz GCs: 4.78 ± 0.25 n=14/6 mice; SGCs: 7.45 ± 0.38 n=13/7 mice, *p<0.0001, Kolmogorov-Smirnov Test). While sEPSC amplitude was higher in SGCs (Fig. 1F, amplitude in pA GCs: 17.80 ± 0.18 n=14/6 mice; SGCs: 19.21 ± 0.26 n=13/7 mice, *p=0.0104, Kolmogorov-Smirnov test), the effect size was small. Similarly, sEPSC 10-90% rise time and biexponential fit to decay were also different between GCs and SGCs, although the effect size was small (Fig. 1G, H, rise time in ms, GCs: 2.13 ± 0.04 n = 14; SGCs: 1.87 ± 0.04 n = 13, *p<0.0001, Kolmogorov-Smirnov test; decay time in ms, GCs: 4.38 ± 0.09 n = 14, SGCs: 3.78 ± 0.11 n = 13, p<0.0001, Kolmogorov-Smirnov test). However, charge transfer was not different between cell types. (Fig. 1I, charge transfer in pA • ms, GCs: 63.71 ± 2.49 n = 14; SGCs: 69.59 ± 2.66 n = 13, p=0.2840, Kolmogorov-Smirnov test). Taken together with our prior finding that SGCs receive more inhibitory synaptic inputs than GCs (Gupta et al., 2012; Afrasiabi et al., 2022), these results indicate that SGCs receive more spontaneous synaptic inputs than GCs.

**Figure 1:**
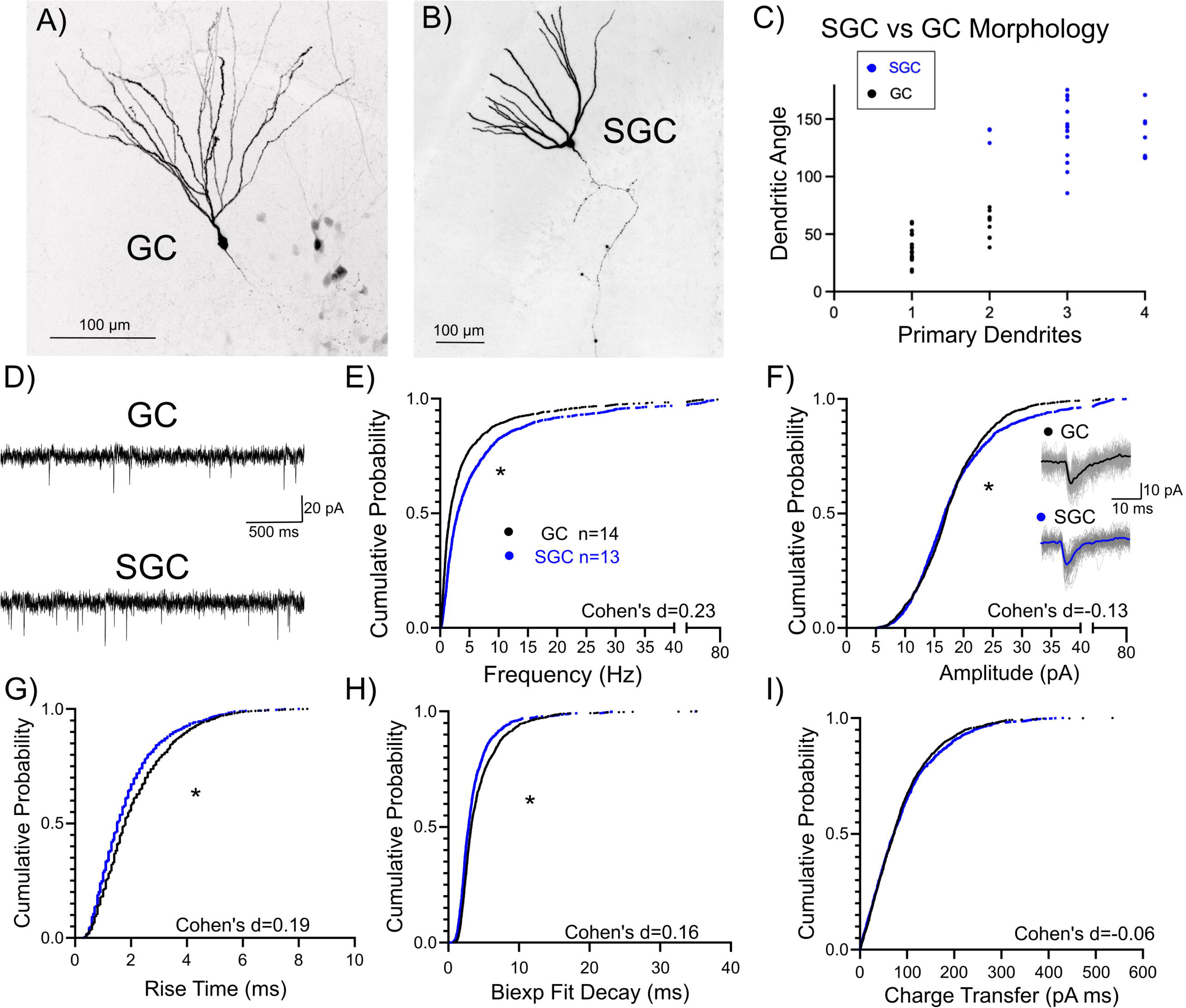
SGCs receive more frequent spontaneous excitatory inputs than GCs. A-B) Representative images of a biocytin-filled GC with a narrow dendritic span and a single primary dendrite (A) and a SGC with a wide dendritic span and multiple primary dendrites (B). Maximum-intensity projections of confocal image stacks are presented as gray-scale, inverted images to visualize hilar axon collaterals. C) Summary plot of dendritic angle and primary dendrites of cells classified as SGC (blue) and GC (black) by the experimenter. D) Representative current traces illustrate spontaneous EPSCs in a GC (top) and an SGC (bottom). E-I) Cumulative probability plot of sEPSC instantaneous frequency (E), amplitude (F), rise time (G), decay (H) and charge transfer (I) in GCs (black) and SGCs (blue). Insets in F: Example of average sEPSC in a representative GC (black) and SGC (blue) overlaid on individual traces in gray. * indicates p<0.05 by Kolmogorov-Smirnov test, n=13-14 cells/group.

### SGCs receive stronger medial entorhinal cortex inputs than GCs

The PP is the primary source of cortical inputs to GCs and SGCs. Consistent with previous findings in rats (Williams et al., 2007)., EPSC amplitude evoked by mass stimulation of the perforant path was not different between GCs and SGCs (Fig 2 A-C, SGC n=6 GC n=7 p=0.68, two-way ANOVA). Because the perforant path includes two distinct input streams which can be differentially recruited based on variations in slice angle, we examined the pathway-specific inputs to GCs and SGCs by selectively expressing the excitatory opsin, channelrhodopsin (ChR2), in the lateral or medial perforant path projections in slices from mice injected with AAV5-hSyn-hChR2(H134R)-EYFP into the lateral or medial entorhinal cortex (Supplementary Figure 1). Dual recordings from SGC-GC pairs, with somata located within 200 µm of each other and at similar depths, were conducted to limit potential confounds due to differences in ChR2 expression between animals and slices (Fig 3 A-C).

**Figure 2:**
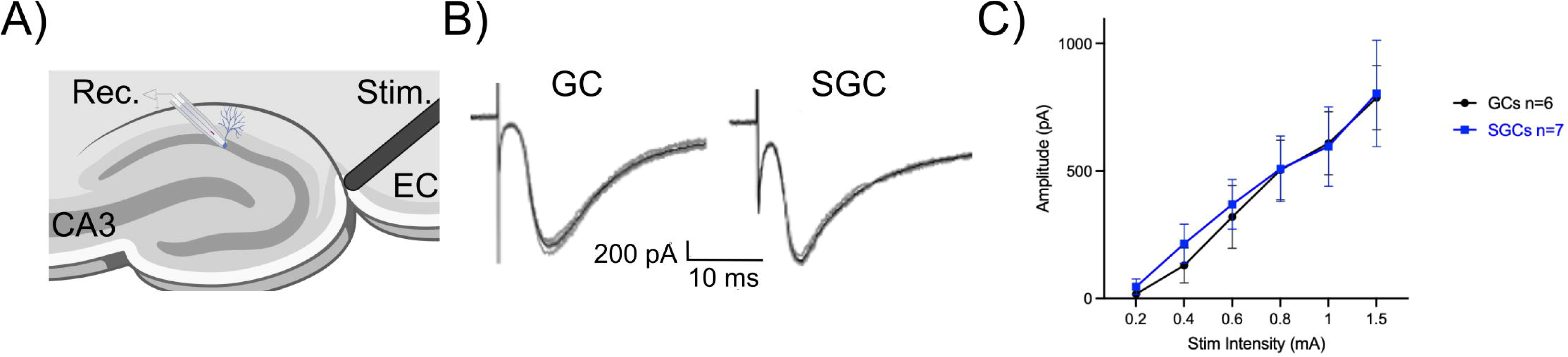
Stimulus-evoked perforant path responses do not differ between cell types. A) Schematic depicting electrode placement for perforant path stimulation. Note that the stimulation electrode was on the other side of the fissure from the outer molecular layer. B) Representative PP-evoked EPSCs in a GC (left) and an SGC (right) at 1.5 mA stimulus intensity. Individual traces are in grey, with the average trace in black. C) Summary plot of perforant path-evoked EPSC peak amplitude in SGCs (blue) and GCs (black) in response to increasing stimulus intensities. n=6-7 cells/group by two-way ANOVA.

**Figure 3:**
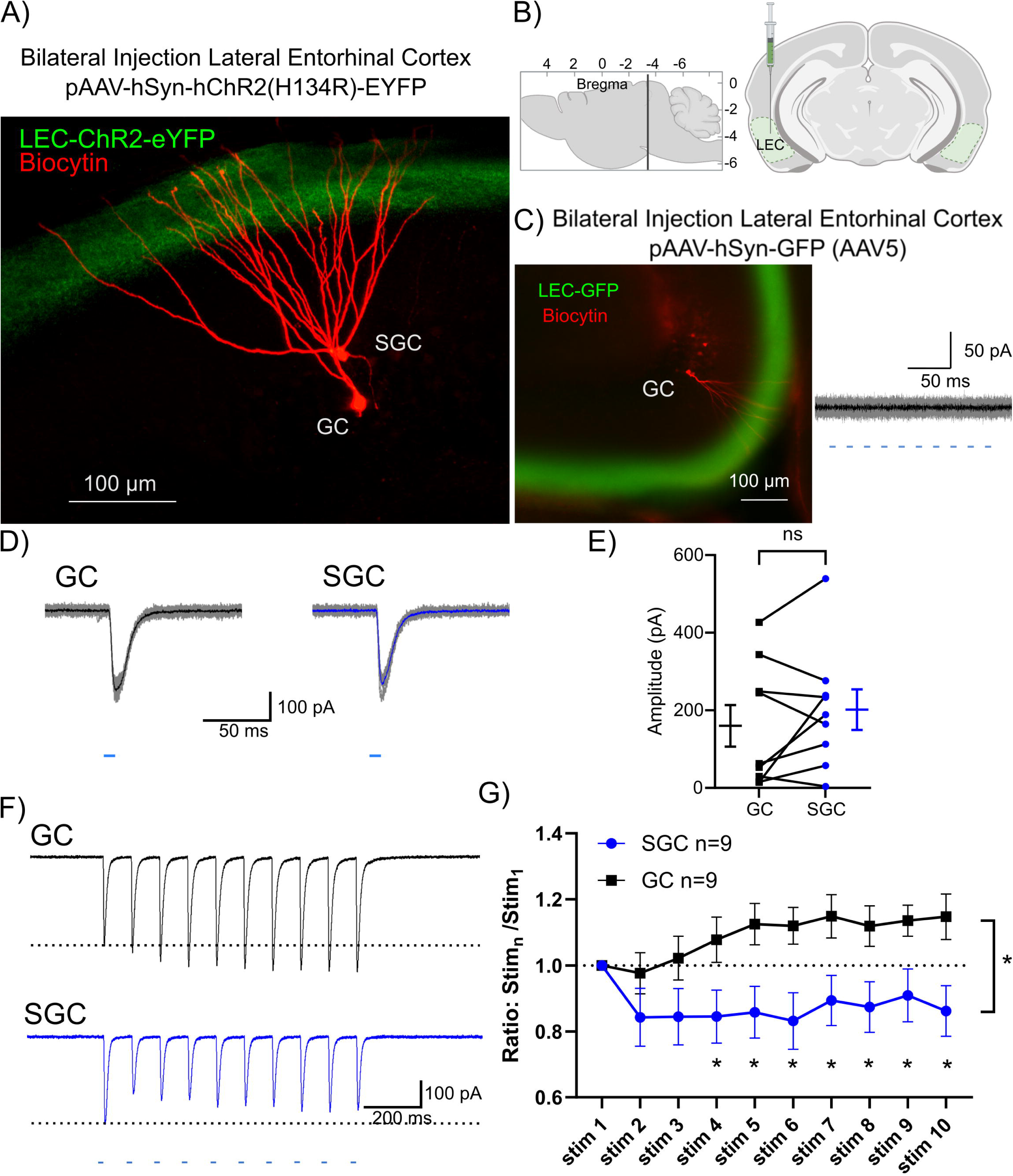
Differential short-term dynamics of LPP inputs to SGCs and GCs. A) Representative image of biocytin-filled SGC and GC (red) labeled during dual recordings. Lateral perforant path projections are labeled with ChR2-eYFP (green). B) Schematic of injection site for virally mediated labeling of projections from the LEC. C) Representative section from a mouse injected with a control virus, pAAV-hSyn-GFP in the LEC to label the LPP (left). The biocytin labeled GC (red) showed no response to blue light pulses (right). D) Representative traces of EPSCs evoked by a single 10 ms light pulse (shown below) in a GC and an SGC. Individual traces are in gray with average trace overlaid in black (GC) or blue (SGC). E) Summary plot of eEPSC peak amplitude in 9 SGC-GC pairs. Mean ± SEM are plotted adjacent to raw data. F) Representative traces of EPSCs evoked by a train of 10 light pulses at 10 Hz in GC (top) and SGC (bottom). Dotted line represents the peak amplitude of the first EPSC. G) Summary plot of the ratio of amplitude of n^th^ evoked EPSC compared to the first EPSC evoked during a 10-pulse train. * indicates p<0.05, for main factor cell type by two-way ANOVA.

As illustrated by an example image of a biocytin-filled GC-SGC pair, AAV injection in the LEC reliably labeled lateral perforant path (LPP) fibers in the OML with ChR2-eYFP (Fig 3A, B). In control experiments, blue light stimulation failed to elicit synaptic responses when mice were injected with control virus (AAV5-hSyn-GFP) lacking ChR2 (Fig. 3B-C). Blue light activation of LEC projections in the OML (0.9 mW, λ=470 nm, 10 ms pulse) evoked low-amplitude EPSCs with no systematic difference between cell types (Fig 3D-E; amplitude in pA, GCs: 160.1 ± 52.66; SGCs: 201.8 ± 51.51; p=0.36, n=9 SGC-GC pairs, Wilcoxon matched-pairs signed-rank test). EPSC latency from the start of optical stimulation was not different between cell types (latency in ms, SGC: 2.45 ± 0.14; GC: 2.43 ± 0.14, p>0.05, Mann-Whitney test), consistent with monosynaptic events. However, the short-term dynamics of the responses to LEC activation at 10 Hz (10 pulses) differed between cell types. While LPP-evoked EPSCs in GCs showed multi-pulse facilitation, EPSCs in SGCs underwent synaptic depression during the same stimulus train (Fig. 3F, G, *p=0.03, two-way ANOVA). Together, these data show that while LPP input strengths are not different between GCs and SGCs, short-term plasticity differs by cell type, which can impact dendritic processing of LEC inputs and activity-dependent recruitment of SGCs and GCs by LEC inputs.

Next, we examined GC-SGC pairs in mice injected with AAV in the MEC to express ChR2-eYFP in medial perforant path fibers in the MML (Fig. 4A, B). As with LPP evoked EPSCs, latency to MPP evoked EPSCs were consistent with monosynaptic events and was not different between cell types (latency in ms, SGC: 2.22 ± 0.33; GC: 2.00 ± 0.14, p>0.05, Mann-Whitney test) In contrast to LEC inputs, optical activation of MEC projections consistently evoked higher amplitude EPSCs in SGCs compared to neighboring GCs in the same slice (Fig. 4C, D, amplitude in pA GCs: 346.2 ± 130.5; SGCs: 609.5 ± 226.0; p=0.016, n=8 SGC-GC pairs, Wilcoxon matched-pairs signed-rank test). Taken together, the stronger MPP synaptic inputs and multi-pulse depression (Fig. 4E, F, p=0.18, two-way ANOVA) of both LPP and MPP synapses observed in SGCs are well suited to recruit SGCs during behaviorally relevant MEC activity.

**Figure 4:**
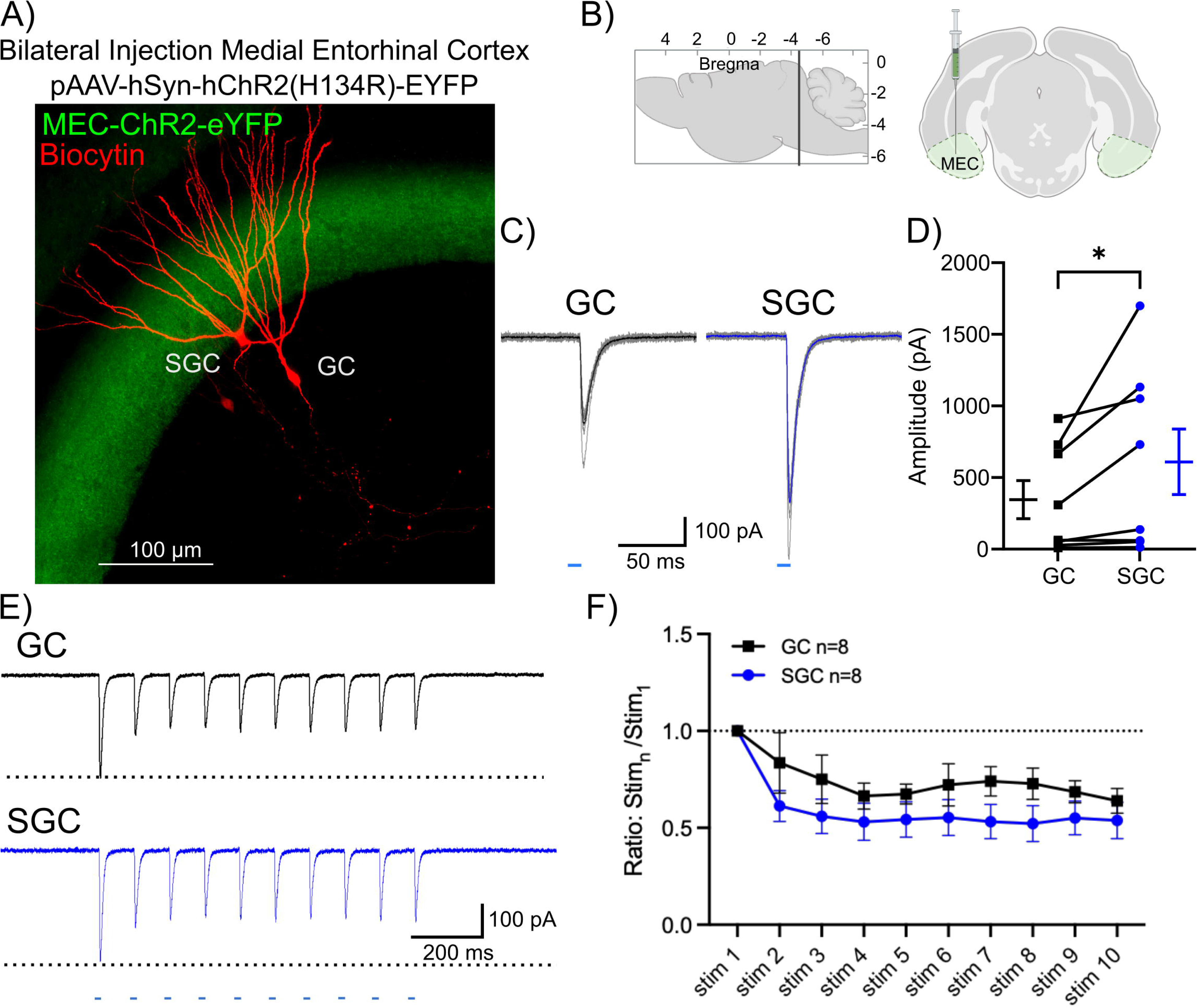
SGCs receive stronger MEC inputs compared to GCs. A) Representative image of biocytin-filled SGC-GC pair (red) in a mouse with AAV injection in the MEC. MEC projections are labeled with ChR2-eYFP(green). B) Schematic of injection sites for virally mediated labeling of projections from the MEC. C) Representative traces of EPSCs evoked by a single 10 ms light pulse in GC and SGC. D) Summary plot of eEPSC peak amplitude in SGC-GC pairs. Mean ± SEM are plotted adjacent to raw data. E) Representative EPSC traces in a GC (top) and an SGC (bottom) evoked by a train of 10 stimuli, 10 Hz light pulses to activate MEC projections. F) Summary plot of the ratio of the amplitude of n^th^ evoked EPSCs compared to the first EPSC evoked during a 10 pulse train. * Indicates p<0.05 by Wilcoxon matched-pairs signed-rank test in n=8 SGC-GC pairs.

### Cell-type specific differences in associational but not commissural inputs between SGCs and GCs

Apart from the perforant path, mossy cells are a major source of glutamatergic commissural and associational inputs to the DG (Buckmaster et al., 1996). Local axon collaterals of mossy cells provide sparse innervation of hilar neurons and local GCs (Botterill et al., 2021). In contrast, long-range mossy cell axons form associational and commissural projections that innervate ipsilateral and contralateral GCs, respectively, along the entire septotemporal axis of the DG (Frotscher et al., 1991; Buckmaster et al., 1996). Consistent with the previous reports in rats (Williams et al., 2007), our slice recordings in mice revealed that the amplitude of hilar stimulation-evoked EPSCs is greater in SGCs than in GCs (Fig. 5, *p=0.01, two-way ANOVA), indicating stronger associational projections to SGCs.

**Figure 5:**
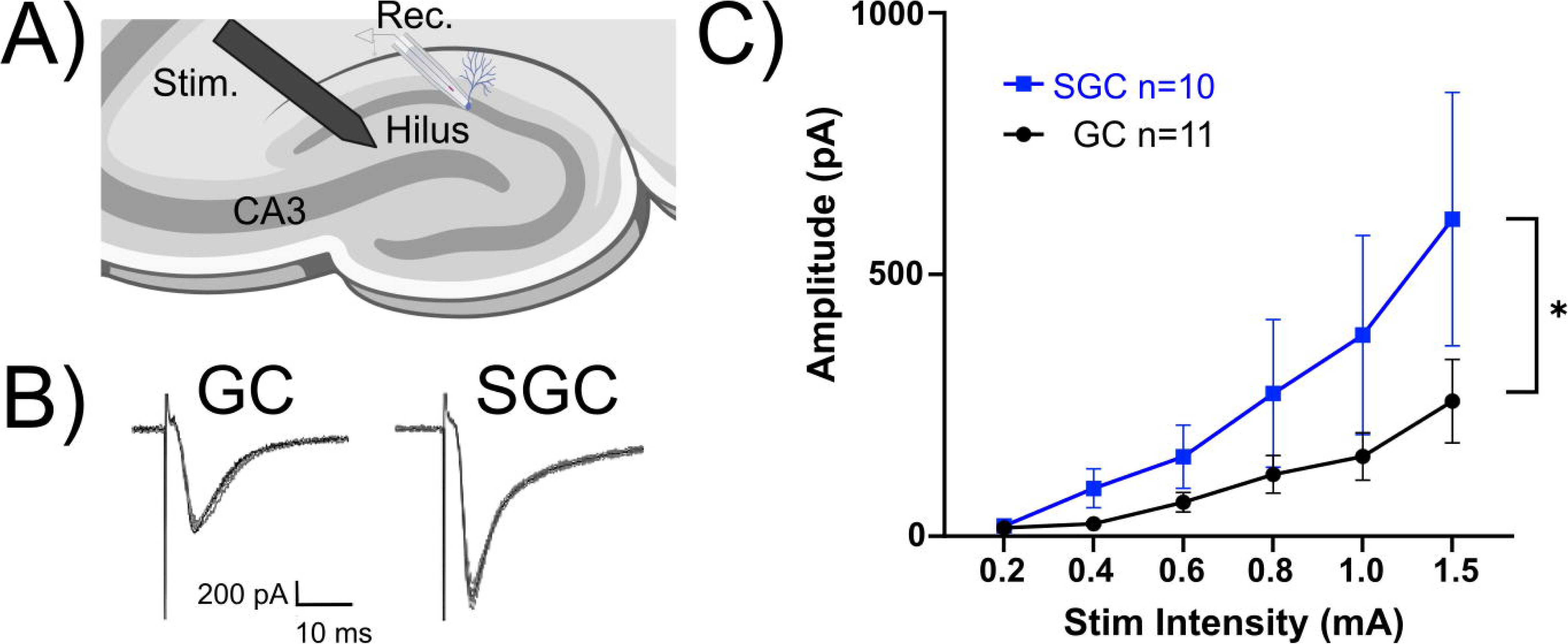
SGCs receive stronger associational synaptic inputs than GCs. A) Schematic depicting electrode placement for hilar stimulation. B) Representative EPSCs in a GC (left) and SGC (right) in response to electrical stimulation in the hilus. Individual traces are in gray, with average trace overlaid in black. C) Summary plot of hilar stimulus-evoked EPSC peak amplitudes at increasing intensities n=5-6 cells/group. * Indicates p<0.05 by two-way repeated-measures ANOVA.

Previous studies have demonstrated that the spatial distribution of mossy cell axon collaterals differs by soma location along the dorsoventral axis of the DG (Houser et al., 2020; Botterill et al., 2021). To independently evaluate ventral and dorsal commissural inputs to GCs and SGCs, we injected AAV5-CAMKIIa-hChR2(H134R)-eYFP unilaterally into the dorsal or ventral hilus to selectively label dorsal and ventral glutamatergic commissural inputs to the contralateral DG. AAV injection in the dorsal hilus consistently labeled diffuse commissural projections extending through the inner and middle molecular layers in the contralateral hilus (Fig. 6A). Optical activation of the dorsal commissural inputs evoked EPSCs in both cell types (Fig. 6B). However, the amplitude of the EPSC evoked by dorsal commissural activation was not different between cell types (Fig. 6C, amplitude in pA GCs: 27.48 ± 5.18; SGCs: 38.07 ± 8.03; p=0.65, n=9 SGC-GC pairs, Wilcoxon matched-pairs signed-rank test). Similarly, multi-pulse depression in response to activation of dorsal commissural inputs at 10Hz (10 stimuli each with 10 ms duration) was not different between cell types (Fig. 6 D, E, p=0.42, two-way ANOVA).

**Figure 6:**
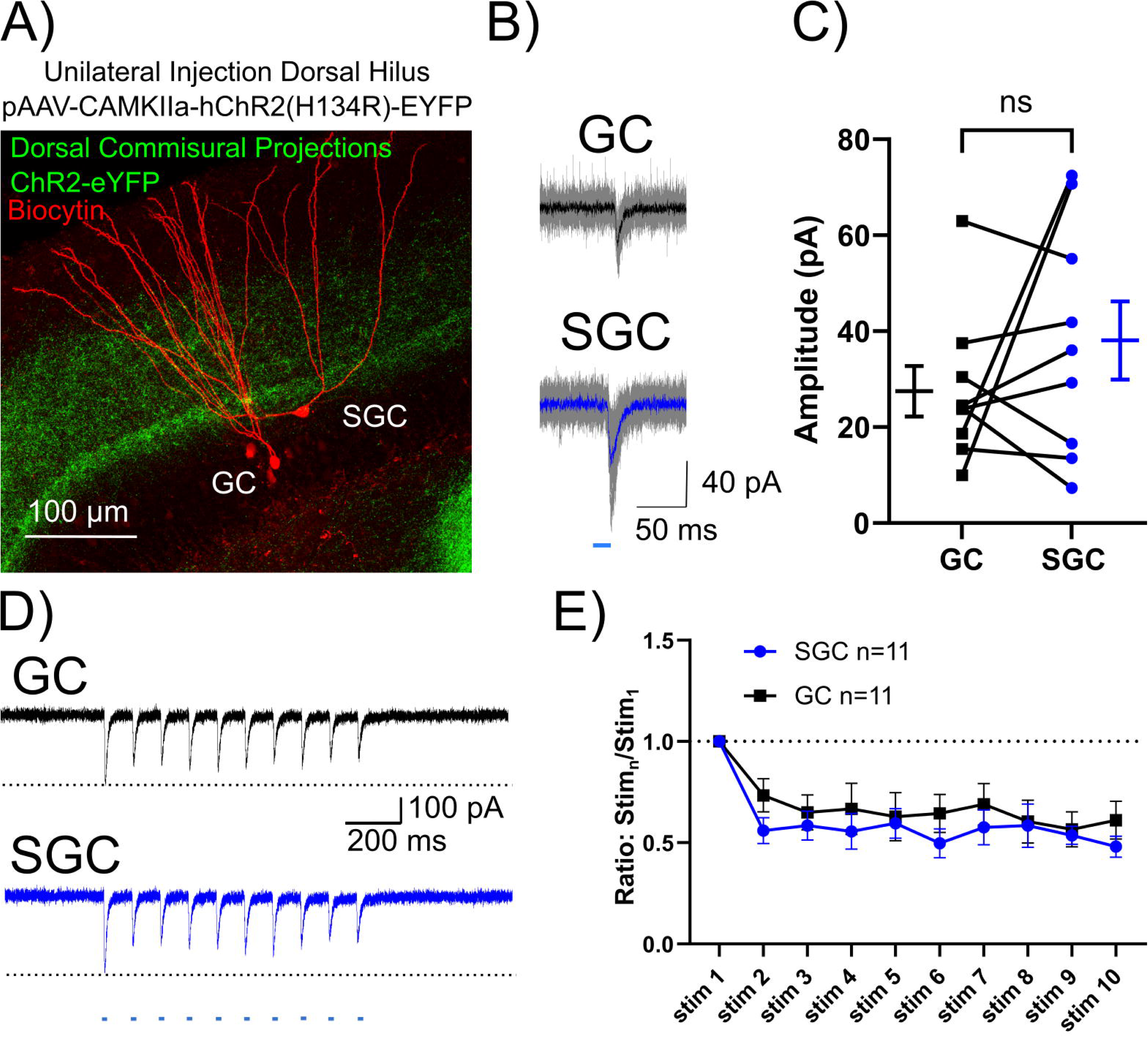
Dorsal commissural synaptic inputs are not different between SGCs and GCs. A) Representative image of biocytin-filled SGC and GC (red) with AAV-driven ChR2-eYFP labeling of commissural projections from the contralateral dorsal hilus (green). B) Representative EPSC traces in a simultaneously recorded GC (top)-SGC (bottom) pair evoked by activation of commissural projections from the dorsal hilus by a single 10 ms light pulse. Individual traces are in gray, with average trace overlaid in black (GC) or blue (SGC). C) Summary plot of EPSC peak amplitude in SGC-GC dual recordings. p>0.05 by Wilcoxon matched-pairs signed-rank test, n=11 SGC-GC pairs. Mean ± SEM are plotted adjacent to raw data. D) Representative traces of EPSCs evoked by a 10-Hz train of 10 ms light pulses, in GC (top) and SGC (bottom). E) Summary plot of the ratio of the amplitude of evoked EPSC compared to the first EPSC evoked in a 10-pulse train.

As reported previously (Houser et al., 2020; Botterill et al., 2021) AAV injection in the ventral hilus labeled a distinct band of fibers restricted to the IML in the contralateral hilus (Fig. 7A). As illustrated by the higher magnification image (Fig. 7B), some of our cell fills revealed extensive SGC axons projections in the IML and GCL prior to entering the hilus, a unique feature of SGCs previously reported in rats (Williams et al., 2007). As with dorsal commissural projections, recordings from GC-SGC pairs revealed no differences in EPSC amplitude (Fig 7C, D, amplitude in pA, GCs: 32.54 ± 9.08; SGCs: 35.66 ± 14.07; p>0.99, n=10 SGC-GC pairs, Wilcoxon matched-pairs signed-rank test) or multi-pulse depression (Fig 7E,F, p=0.93, two-way ANOVA) between GCs and SGCs. In control experiments in which mice were injected with AAV5-CAMKIIa-GFP lacking ChR2 in the ventral hilus, optical activation failed to elicit synaptic responses in the recorded GCs (not shown). Collectively, these data demonstrate that while SGCs receive stronger hilar associational-evoked EPSCs than GCs, the commissural inputs from the dorsal and ventral hilus are not different between cell types.

**Figure 7:**
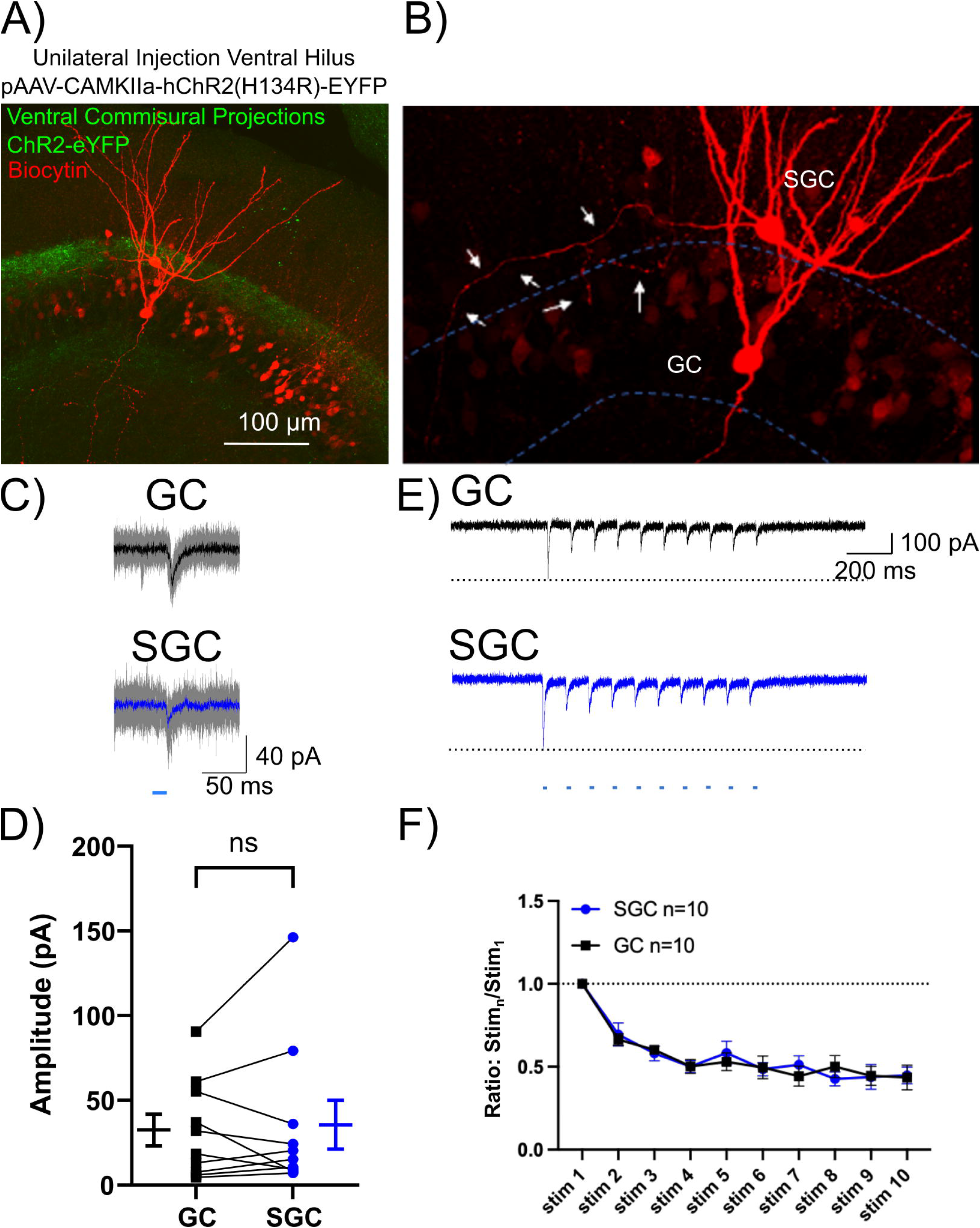
Ventral commissural synaptic inputs are not different between SGCs and GCs. A) Representative image of biocytin-filled SGC and GC (red) with AAV driven ChR2-eYFP labeling of commissural projections from the ventral hilus (green). B) Higher magnification image of the same cell pair in (A) shows biocytin-filled axon traversing the IML and extending collaterals in the GCL denoted by white arrows. C) Representative EPSC traces evoked by a single 10 ms light pulse activation of commissural projections from ventral hilus in GC (top) and SGC (bottom). Individual traces are in gray with average trace overlaid in black (GC) or blue (SGC) in n=10 SGC-GC pairs. D) Summary plot of eEPSC peak amplitude in SGC-GC dual recordings. Mean ± SEM are plotted adjacent to raw data. E) Representative traces of EPSCs evoked by a 10-Hz train of 10 ms light pulses in GC (top) and SGC (bottom). F) Summary plot of the ratio of amplitude of evoked EPSCs compared to the first EPSC evoked in a 10-pulse train.

### Subcortical inputs from the supramammillary nucleus are not different between GCs and SGCs

In addition to entorhinal and commissural inputs, the DG also receives subcortical inputs from the supramammillary nucleus (SuM) in the hypothalamus (Soussi et al., 2010). SuM axon terminals are distributed in the supragranular region of the GCL of the DG and form monosynaptic connections on GCs, which co-release glutamate and GABA (Soussi et al., 2010; Ajibola et al., 2021). To examine SuM inputs to GCs and SGCs, we undertook dual SGC-GC recordings of both EPSCs and IPSCs using a Cs-based internal solution in mice that received bilateral injection of AAV-hSyn-ChR2-eYFP in the SuM (Fig. 8A, B). As in GCs, activation of SuM inputs elicited both IPSCs and EPSCs in SGCs, demonstrating functional SuM inputs to SGCs. However, the amplitude of EPSC in response to photostimulation of SuM terminals was not different between cell types (Fig. 8D, amplitude in pA: GCs: 16.92 ± 4.07; SGCs: 29.25 ± 11.54; n=10 pairs p=0.63, Wilcoxon matched-pairs signed rank test). Similarly, the amplitude of SuM-evoked IPSC was not different between cell types (Fig. 8E, amplitude in pA, GCs: 47.17 ± 7.77, SGCs: 36.76 ± 10.44, n=10 pairs, p=0.63, Wilcoxon matched-pairs signed rank test). These data identify that SGCs receive SuM inputs and that the strength of SuM inputs is not different between GCs and SGCs.

**Figure 8:**
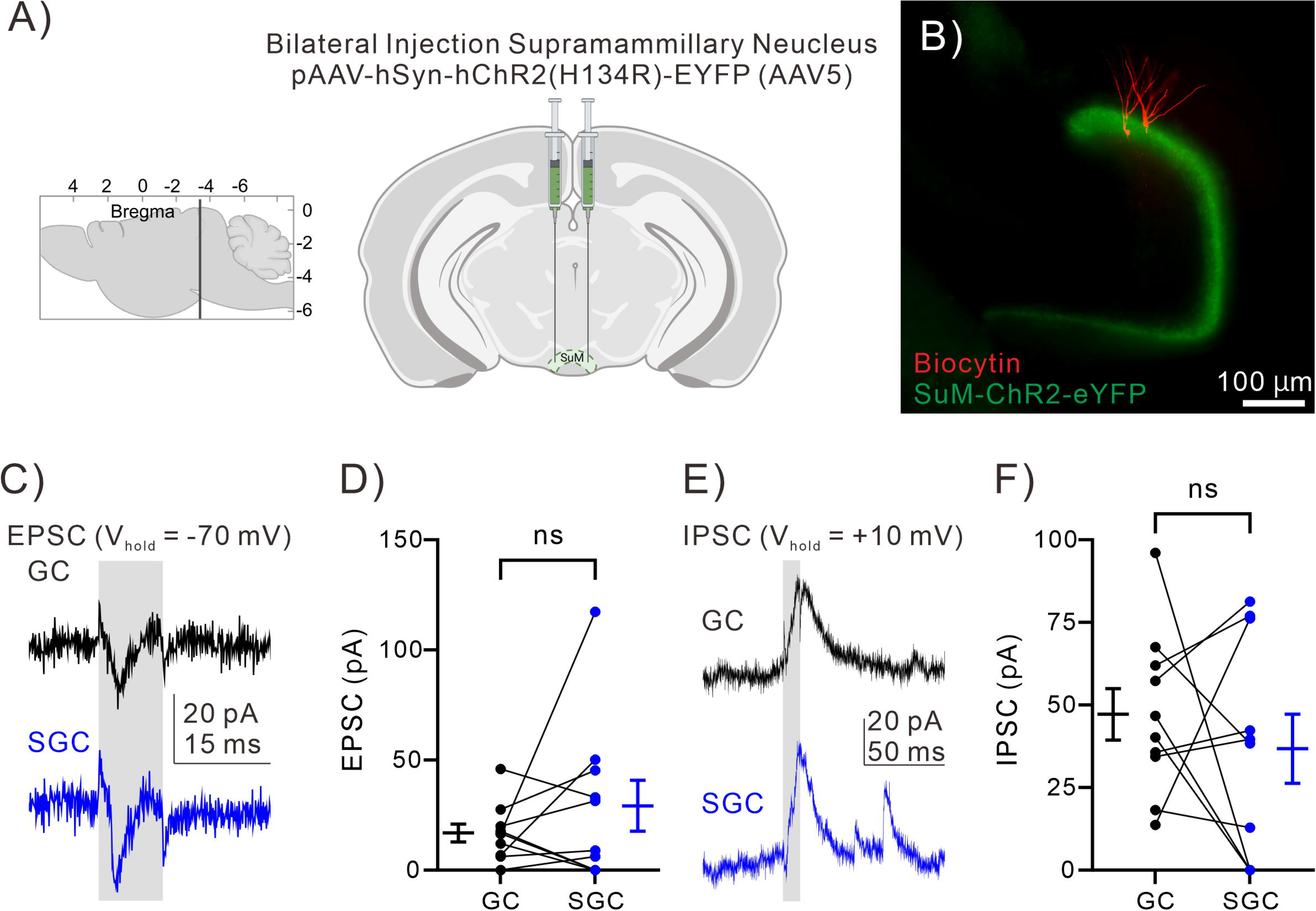
SuM-evoked responses did not differ between CGs and SGCs. A) Schematic of injection sites for virally mediated labeling of projections from the supramammillary nucleus (SuM). B) Representative image of biocytin-filled SGC and GC (red). SuM inputs are labeled with ChR2-eYFP (green). C) Photostimulation-evoked EPSCs in a GC (top) and an SGC (bottom) under voltage-clamp configuration at a holding potential of -70 mV. The grey box illustrates photostimulation. D) Summary plot of EPSC peak amplitude in SGC-GC pairs (n=10 SGC-GC pairs). Mean ± SEM are plotted adjacent to the raw data. E) Photostimulation-evoked IPSCs in a GC (top) and an SGC (bottom) under voltage-clamp configuration at a holding potential of +10 mV. The grey box illustrates photostimulation. F) Summary plot of IPSC peak amplitude in SGC-GC pairs (n=10 SGC-GC pairs). Mean ± SEM are plotted adjacent to the raw data.

### Dendritic spine density is higher in SGCs than in GCs

Given the higher sEPSC frequency and stronger MEC inputs, we examined dendritic spine density in biocytin-filled SGCs and GCs obtained during physiological recordings. Images were captured from 3 dendritic segments: proximal, middle, and distal to the soma (illustrated in Fig 9A-B). Remarkably, on occasion we found spines on both somata and on putative SGC axons (Fig 9C), identified based on the recovery of the extensive axonal arbor in the hilus. These putative axonal spines appear well positioned to receive associational inputs in the IML. However, none of the GCs imaged as part of this study exhibited somatic or axonal spines. Dendritic spine reconstructions revealed higher overall spine density in SGCs than in GCs (Fig 9D; Spines / µm: SGCs: 1.12 ± 0.08, GCs: 0.81 ± 0.07, *p=0.013 by nested t-test. n=6 cells/group). Evaluation of spine density relative to distance from soma identified that the spine density in the middle and proximal dendritic segments of SGCs was significantly higher than in GCs (Fig. 9A-B, E, proximal spine density in Spines / µm, GCs: 0.50 ± 0.03, n = 5, SGCs: 1.04 ± 0.11, n = 6, *p=0.001; middle spine density, GCs: 0.80 ± 0.08, n = 6, SGCs: 1.24 ± 0.12, n = 6, *p=0.0034, two-way ANOVA). Note that because of their wide dendritic arbor and somatic location in the IML, a majority of SGC proximal and middle dendrites are in the MML, which receives MPP inputs. However, spine density in the distal dendritic segments in the OML innervated by the LPP did not differ between cell types (Fig. 9A-B, E, distal spine density in Spines / µm, GCs: 0.96 ± 0.13, n = 6, SGCs: 1.08 ± 0.09, n = 6, p=0.408, two-way ANOVA).

**Figure 9:**
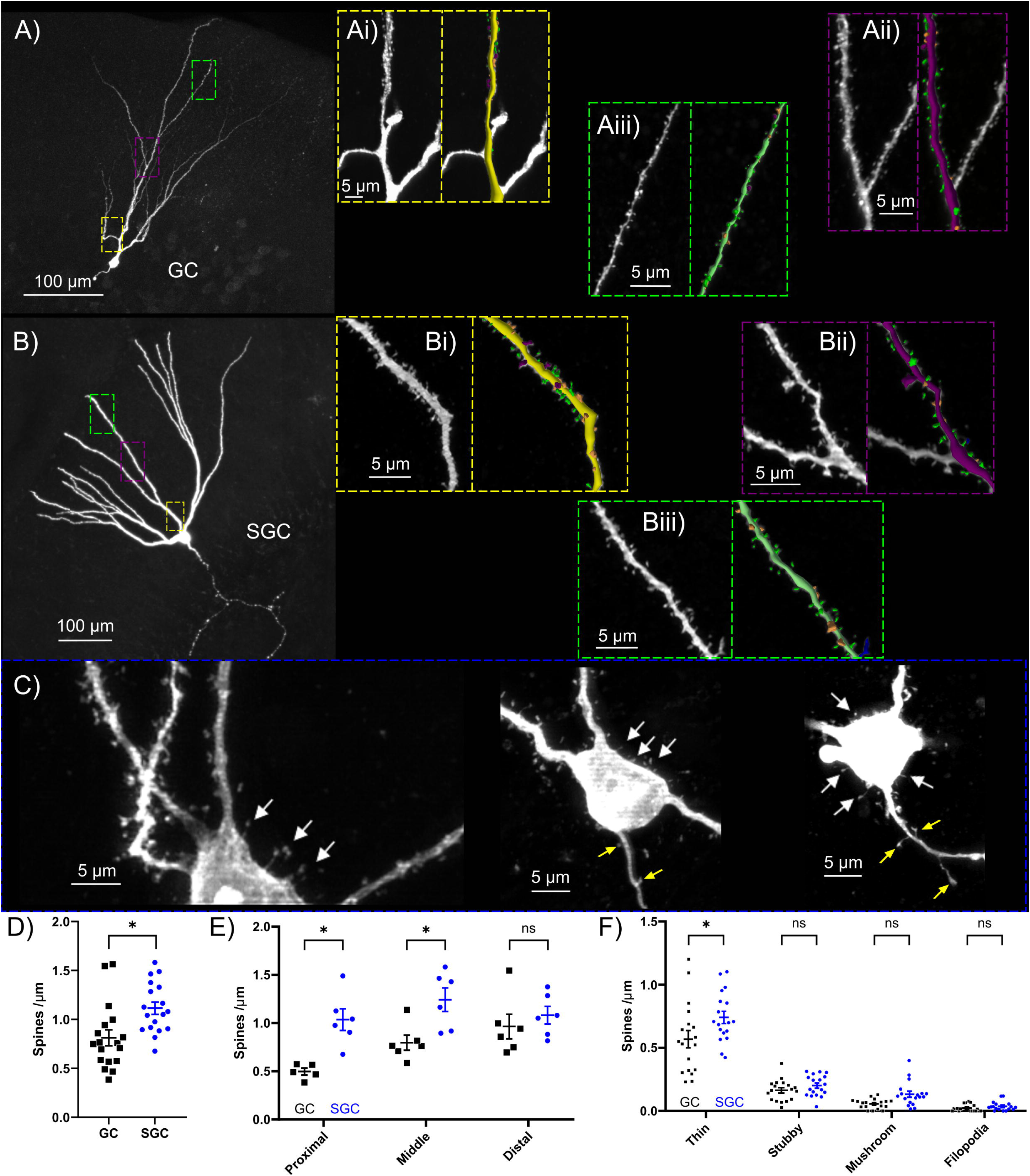
SGCs have higher dendritic spine density than GCs. A-B) Representative confocal images of a GC (A) and an SGC (B) with insets illustrating reconstructed spines in proximal (yellow), middle (purple), and distal (green) dendritic segments. C) Representative images of 3 SGC somata demonstrating the presence of somatic spines (white arrows) as well as axonal spines (yellow arrows). D) Summary plot compares spine density between cell types. * Indicates p<0.05 by Mann Whitney n=18 segments/6cells/group. E) Summary plots of layer-specific differences in spine densities between cell types. F) Summary plots of of thin, stubby, mushroom and filopodial dendritic spines in cell types. * Indicates p<0.05, by two-way ANOVA with Holm-Šídák’s multiple comparisons *post hoc* test in n=6 cells/group.

Spines can be categorized based on their shape, and spine morphology is associated with distinct functional characteristics. We examined thin, stubby, mushroom, and filopodial spine types, classified using automated algorithms in Neurolucida 360 (Rodriguez et al., 2008), followed by validation by the experimenter, to determine whether SGCs preferentially express an excess of a specific dendritic spine subtype. Thin spines, which can be transient and are referred to as “learning spines” due to their proposed involvement in plasticity (Popov et al., 2004; Bourne and Harris, 2007), were more abundant in SGCs compared to GCs (Fig 9F). Further analysis of the density of spine types by dendritic segment in a subset of cells revealed a selective increase in thin and mushroom spines in the middle dendritic segment of SGCs (Supplementary Figure 2, Supplementary Table 1). SGC somatic spines appear to be a mix of filopodial and thin spines. Collectively, these data indicate that SGCs with somatic spines and increased proximal and middle dendritic spines are well positioned to receive more associational and MPP inputs.

### Differential recruitment of GCs and SGCs by LEC and MEC inputs

Because LEC and MEC inputs showed pathway-specific differences in input strength and short-term dynamics, we examined whether they differ in their ability to elicit spiking in GCs and SGCs. Cell-attached recordings from GC-SGC pairs in response to a 10-ms photostimulation of LPP (75% light intensity, 35 mW) failed to identify a systematic difference in firing probability between GCs and SGCs (Fig. 10A, B, probability in % GCs: 45 ± 20.62; SGCs: 83.33 ± 16.67; p=0.25, n=6 SGC-GC pairs, Wilcoxon matched-pairs signed-rank test), consistent with the lack of difference in LPP-evoked EPSC amplitude between the cell types. To examine short-term dynamics, we used 10 Hz LPP stimulation at 50% of the intensity that evoked maximum firing. As a result, LPP inputs to SGCs and GCs were activated at different light intensities. Consistent with the facilitation of EPSCs, GC firing probability increased in response to 10-Hz stimulation of LPP (Fig. 10C, D). In contrast, SGC firing probability trended to decline during 10-Hz photosimulation of LPP inputs (Fig. 10C, E), consistent with short-term depression of LPP inputs to SGCs.

**Figure 10:**
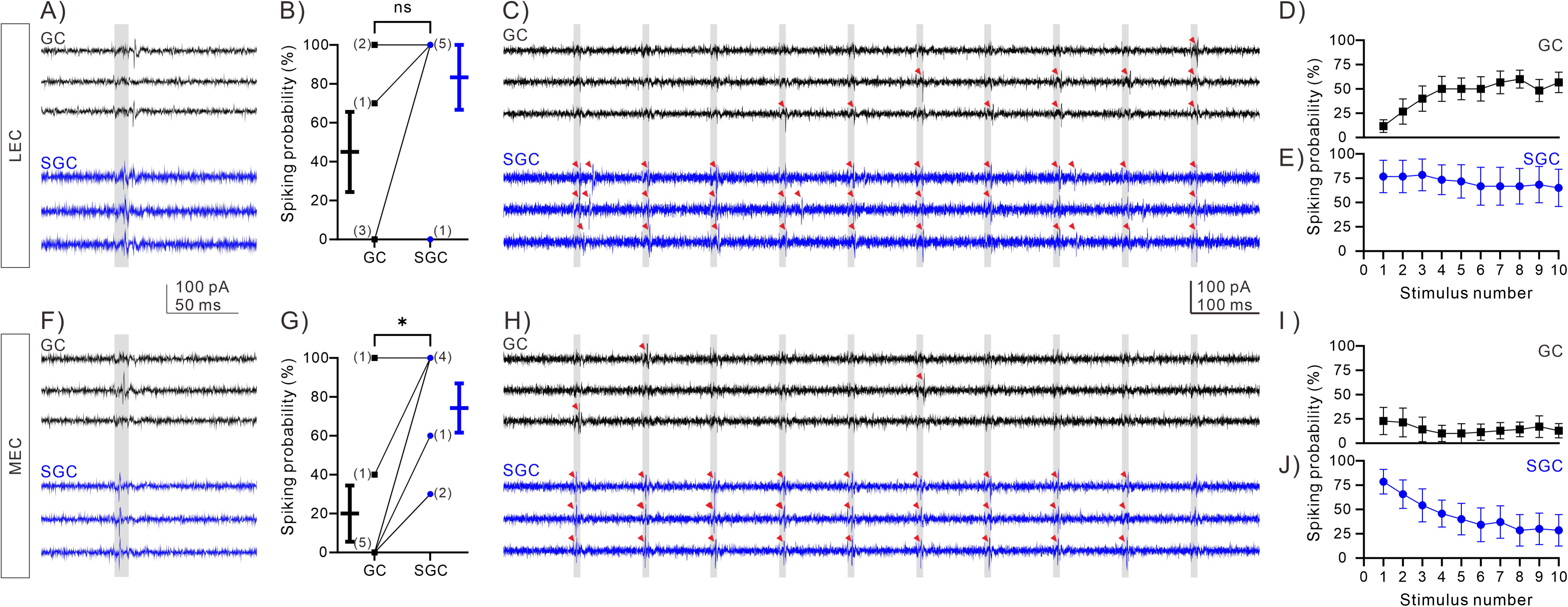
Differential recruitment of GCs and SGCs by LEC and MEC inputs. A) Membrane responses to a 10-ms photostimulation (grey box) of LPP (75 % light intensity) in an SGC-GC pair recorded in cell-attached configuration. B) Spiking probability of each SGC-GC pair (n=6 SGC-GC pairs). Note that the counts of overlapping data points are presented in parentheses. Mean ± SEM are plotted adjacent to the raw data. C) Spikes (red arrowheads) in a representative cell-attached recording from a GC (black) and an SGC (blue) during 10-Hz photostimulation at LEC inputs. Grey boxes indicate photostimulation. D-E) Summary plots of spike probability in GCs (D) and SGCs (E) in during 10-Hz photostimulation of LEC inputs. F) Membrane responses to a 10-ms photostimulation (grey box) of MPP (75 % light intensity) in an SGC-GC pair recorded in cell-attached configuration. G) Spiking probability of each SGC-GC pair (n=7 SGC-GC pairs) during MPP stimulation. Note that the counts of overlapping data points are presented in parentheses. Mean ± SEM are plotted adjacent to the raw data. H) Representative cell-attached recordings illustrate spikes (red arrowheads) in a GC (black) and an SGC (blue) during 10-Hz photostimulation of MEC inputs. Grey boxes illustrate photostimulation. I-J) Summary plots of spike probability in GCs (I) and SGCs (J) during 10-Hz photostimulation of MEC inputs.

Consistent with MPP activation evoking higher EPSC amplitude in SGCs, the probability of SGC firing in response to photostimulation of MPP (75% light intensity, 35 mW) was consistently greater than in GCs (Fig. 10F, G, probability in % GCs: 20 ± 14.47; SGCs: 74.29 ± 12.70; *p=0.0312, n=7 SGC-GC pairs, Wilcoxon matched-pairs signed-rank test). In experiments evaluating short-term plasticity, both SGCs and GCs showed a decline in firing probability during 10-Hz photosimulation of MPP inputs (Fig. 10H,I, J), consistent with short-term depression of MPP-evoked EPSCs in both GCs and SGCs. Note that because SGCs and GCs were activated at different light intensities, we do not directly compare the rate of short-term dynamics between the cell types. Importantly, these results show that MPP inputs recruit SGCs more reliably than GCs.

### Cell morphology does not impose differences in dendritic EPSP integration between GCs and SGCs

Given the layer-specific inputs and the potential role of passive dendritic structures in regulating synaptic signal processing, we implemented morphologically constrained multicompartmental models of GCs and SGCs to evaluate whether neuronal structure contributes to differences in dendritic attenuation and synaptic integration in GCs and SGCs. Accurate 3D reconstructions of 5 biocytin-filled GCs and SGCs, generated using Neurolucida 360, were exported into “Trees” software (Cuntz et al., 2011) to segment dendrites into functionally distinct thirds representing the inner, middle, and outer molecular layer synaptic input zones. Segmented GC/SGC morphologies were imported into NEURON (Hines and Carnevale, 1997) to implement active and passive conductances for functional simulations (Santhakumar et al., 2005). Channel densities, including leak conductance, were kept identical across all neurons to isolate the impact of neuronal morphology. Consistent with the difference in soma shape between GC and SGC, we modeled the SGC somatic compartment as larger than in GCs. The resting membrane potential (RMP) of model cells was set at -70 mV and was not different between model and biological cells (RMP in mV, model GCs: 70.51 ± 0.14, n=4; biological GCs: 78.25 ± 3.86, n=4; model SGCs: 70.25 ± 0.05, n=4; biological SGCs: 72.76 ± 3.32, n=5, p=0.351 F (1, 13) = 0.94, two-way ANOVA). Similar to biological cells (Williams et al., 2007; Gupta et al., 2012; Afrasiabi et al., 2022; Dovek et al., 2025), input resistance (R_in_) of model SGCs was lower than in the model GCs (R_in_ in MΩ, model GCs 179.4 ± 18.97; model SGCs: 99.95 ± 18.77, n=4; p<0.05, t-test).

Identical biexponential synaptic conductance simulating AMPA synapses (Golding et al., 2005) were located in dendritic segments corresponding to the three major input layers (IML, MML, and OML). The local (dendritic) and somatic voltage changes (EPSPs) in response to activation of the synaptic conductance were measured to evaluate the effect of morphology on dendritic attenuation of EPSP amplitude between cell types (Fig. 11A). Comparison of the local dendritic voltage revealed that the local EPSP evoked by an identical conductance was significantly higher in GC than in SGC regardless of dendritic location (Fig. 11B, Supplementary Table 2). Moreover, the proximal-to-distal boosting of dendritic amplitude, while present, was considerably attenuated in SGCs. The peak amplitude of somatic EPSPs in response to an identical dendritic conductance was significantly higher in GCs than in SGCs (Fig. 11C, Supplementary Table 2). Somatic EPSP amplitude, measured as a percent of the corresponding local dendritic EPSP, was lower when synapses were positioned farther from the soma, consistent with greater attenuation of distal synapses (Fig. 11D, Supplementary Table 2). Despite the significant effect of dendritic distance on input attenuation, cell type did not affect the extent of amplitude attenuation (Fig. 11D, Supplementary Table 2).

**Figure 11:**
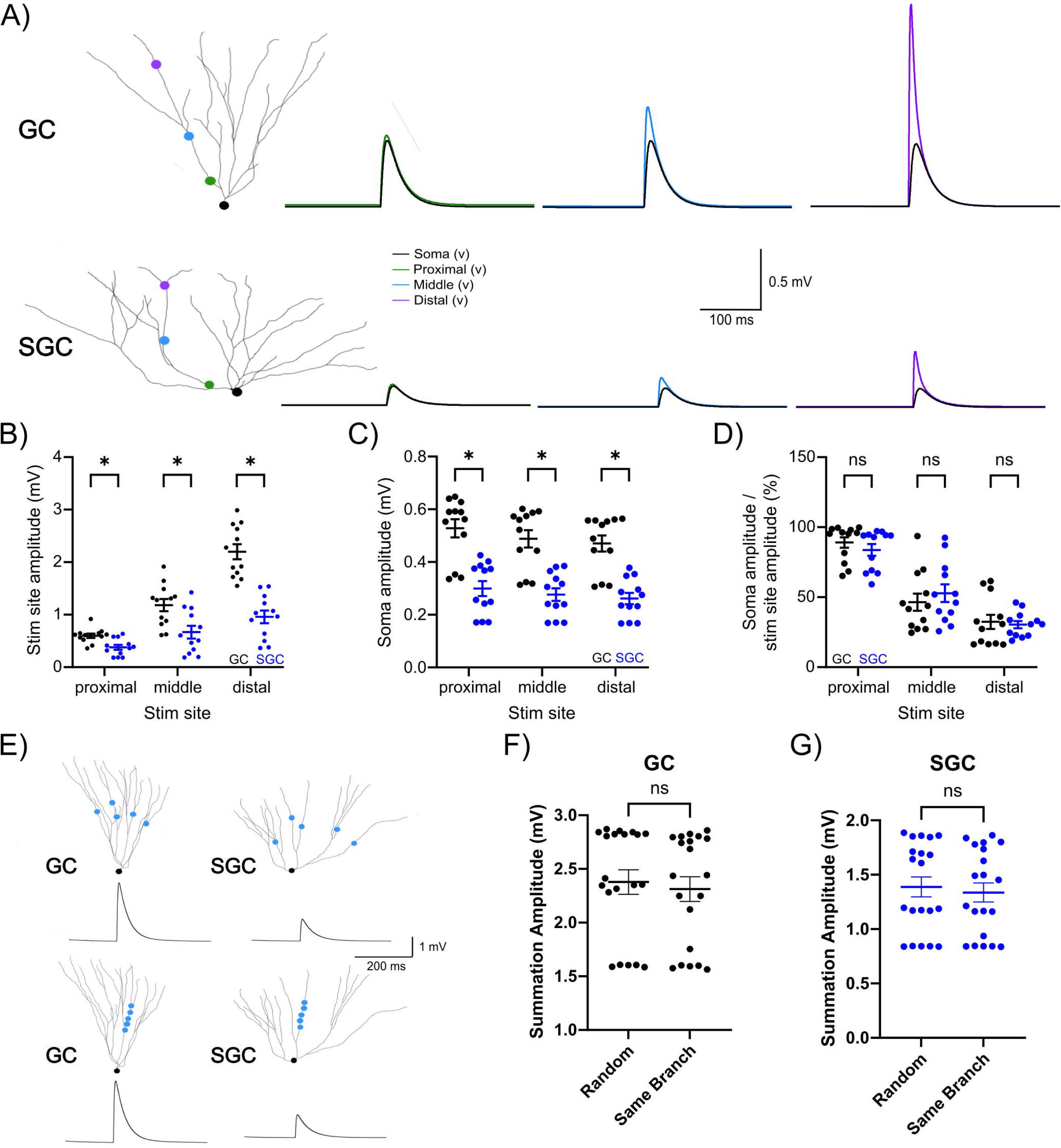
Morphometric computational models reveal greater dendritic boosting of EPSP in GCs than in SGCs. A) Representative reconstructions of a GC (top) and an SGC (bottom), and membrane voltage traces show EPSPs in response to activation of identical AMPA conductance at various dendritic locations. Traces illustrate EPSP amplitude at the soma (black) and at the local input site in the IML (green), MML (blue), and OML (purple). B-C) Amplitude of EPSPs at the site of input on the dendritic segment (B) and at the soma (C). *indicates p<0.05 by two-way ANOVA with Holm-Šídák’s multiple comparisons *post hoc* test in n= 3 trials/4 cells. D) Summary plot of attenuation comparing amplitude at the site of the stimulus to the amplitude recorded at the soma. E) Representative reconstructions of GC (left) and SGC (right) illustrate the location of 5 input synapses in the middle molecular layer on either different branches selected at random (top) or on the same dendritic branch (bottom). Examples of summated EPSPs are illustrated below. F-G) Summary plot of the summated EPSP amplitude in GCs (F) and SGCs (G) when the stimuli were on either the same branch or different branches. n=10 trials/cell, 4 cells/group.

Since we identified stronger MPP inputs to SGC, we focused on the middle dendritic segment and examined the summation of inputs to the same or different dendritic branches (Fig. 11E). We distributed five identical AMPA synaptic conductances either within a single middle dendritic segment on the same branch (5 trials/cell) or randomly in middle dendritic segments across dendritic branches (5 trials/cell). The amplitude of the summed EPSP at the soma resulting from simultaneous activation of all 5 synapses was consistently lower than the numerical sum of the somatic EPSPs in response to activation of the individual synapses, indicating sublinear input summation in both cell types. Moreover, synaptic input summation did not differ whether the same synapses were distributed across 5 randomly selected branches (random) or located within the same dendritic branch (same branch) in either cell type (Fig. 11F, G). Overall, the simulations indicate that dendritic structure does not differentially impact synaptic input attenuation or summation in GCs and SGCs.

### Preferential recruitment of SGCs to DG neuronal ensembles is independent of the behavioral task

Since we find that SGCs have a greater basal excitatory drive and are more reliably recruited by MPP input compared to GCs, we adopted TRAP2::tdT transgenic mice to induce c-Fos-driven tdT (reporter) labeling of DG ensembles (DeNardo et al., 2019) in order to evaluate competing hypotheses concerning SGC recruitment to active neuronal ensembles. Specifically, we tested whether more frequent spontaneous glutamatergic drive and sustained firing made SGCs more active regardless of task, or whether SGCs were selectively activated by the MPP contextual input stream. Although task-related activation of SGCs has been reported previously (Erwin et al., 2020; Dovek et al., 2025; Dovek et al., 2026), how this recruitment compares with basal c-*Fos*-driven labeling of SGCs in task-naïve mice housed in their home cage remains to be evaluated. Since SGCs represent approximately 5% of the GC population (Save et al., 2019), we reasoned that a greater than 5% reporter labeling in SGCs in control mice housed in the home cage with a reduction in proportional activation during a contextual task would suggest that SGCs have elevated activity levels unrelated to behavioral tasks. Alternatively, a low 5% basal SGC labeling in task-naïve mice with labeling of a greater proportion of SGCs as part of active neuronal ensembles in mice trained in a contextual spatial learning task would suggest selective SGC activation by contextual information. Interestingly, prior studies show that the relative SGC recruitment did not appear to depend on task type (Erwin et al., 2020; Dovek et al., 2025). We used task-naïve controls and littermate mice trained in a DG-dependent Barnes maze task to test whether SGC recruitment depends on active neuronal ensembles. Experimental mice were trained for 6 days on the Barnes maze to use spatial cues to find an escape hole, followed by a probe trial and 2 days of reacquisition trials (Re-aq) (Fig. 12A). Task-naïve littermates were brought to the testing room but were never exposed to the Barnes maze table. Evaluation of primary latency, primary errors, total latency, and total errors for mice to locate the escape box confirmed the progressive improvements in task performance from acquisition days 1 through 6 (Fig. 12B-E). BUNS analysis (Illouz et al., 2016) assessing the use of spatial search strategy revealed that mice shifted from random or serial search to a spatial strategy as they progressed through acquisition days. A concatenated version of their search strategy combining all strategies that involved some aspect of spatial awareness into a “spatial approach” further confirmed maximal use of the spatial strategy by day 6 (Fig. 12E, F). Additionally, cognitive scores based on the strategy (developed by Illouz *et al*., 2016) demonstrated progressive improvement from days 1 through 6 of acquisition (Fig. 12G, p<0.00001 by Kruskal-Wallis ANOVA with Dunn’s multiple comparisons, n=14 mice). Having established that test animals reliably adopt a spatial search strategy by day 6, we injected 4-hydroxytamoxifen (4-OHT) 10 minutes before day 6 of the acquisition trial to induce tdT expression in neurons expressing the immediate early gene c-*Fos* during behavior testing. Littermate controls in their home cage also received 4-OHT injections and were returned to the home cage. To minimize non-specific activation, test mice and their littermate controls were housed in the test room for 5 hours prior to and following injection and testing, before returning to the vivarium. A probe trial on day 7, during which the escape was removed, showed that mice with acquisition training spent significantly more time in the quadrant in which the escape was previously located compared to untrained littermate control animals (Fig. 12H, time in the escape zone in sec, Control: 17.00 ± 5.04, n=8, Test: 48.37 ± 4.65, n=14, *p=0.0004, Mann-Whitney test), confirming the use of a spatial search strategy in trained animals.

**Figure 12:**
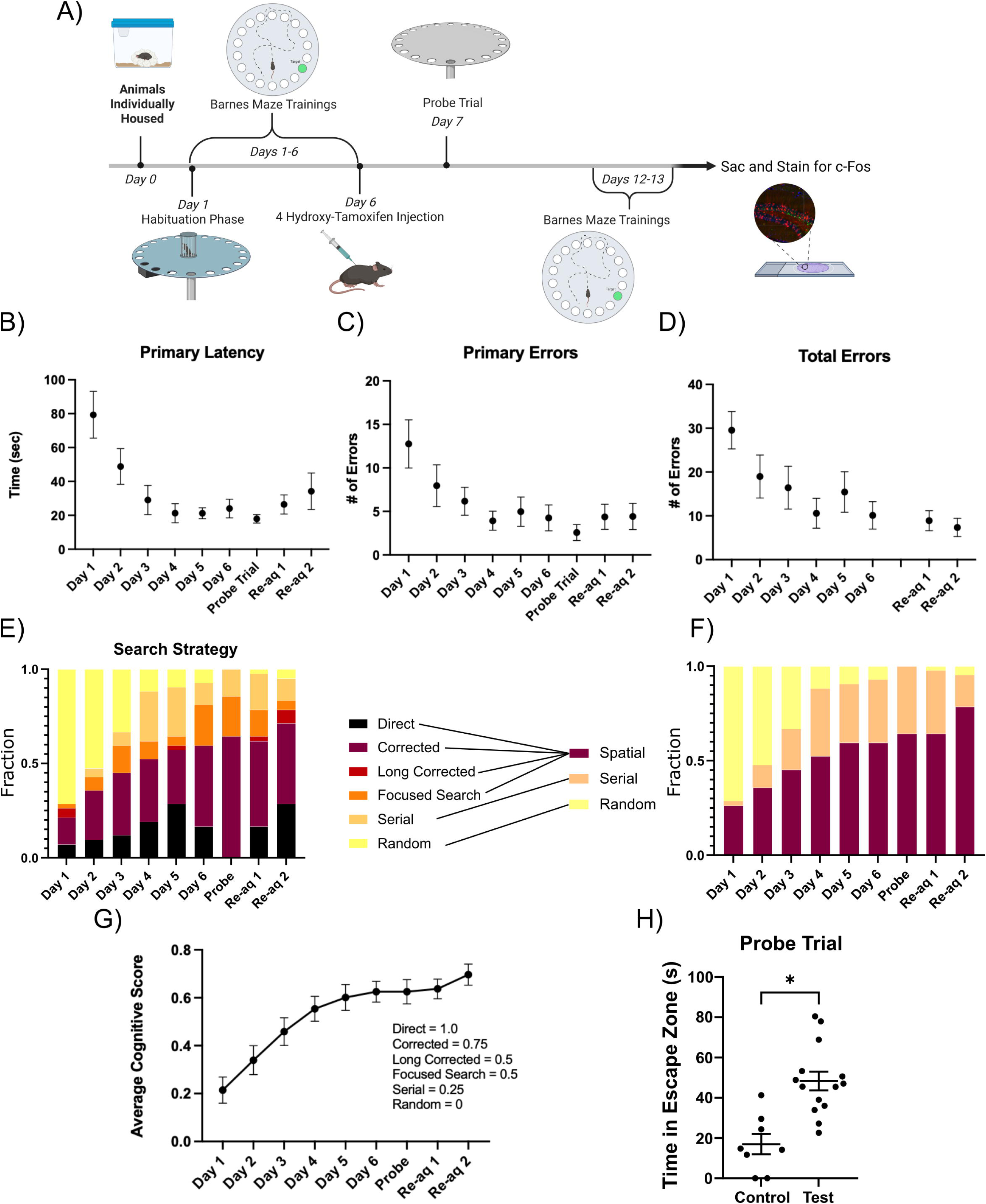
Animals show steady improvement to using a spatial strategy in the Barnes Maze. A) Schematic of experimental timeline for test mice. B-D) Summary plots of primary latency, or the time it takes for the animal to first inspect the escape hole (B), and primary errors, or the number of errors until first exit inspection (C), and total errors (D). E) BUNS output of search strategy analyzed and how we chose to refine it (right) by combining all strategies that use any form of spatial awareness into a single “spatial” category. F) Summary of search strategies used by mice in the Barnes maze task. G) Summary of cognitive scores, derived from search strategy used during task acquisition and probe trials. H) Summary plot of overall time spent in escape quartile during the probe trial. * indicates p<0.05 by Mann-Whitney test.

Having established that, unlike task-naïve controls, mice trained in the Barnes maze reliably adopt a spatial strategy, we quantified the number of neurons labeled with tdT in sections from animals labeled on day 6 of acquisition and task-naïve littermate control animals. While there was considerable variability in the number of tdT-labeled neurons between animals, there was a trend towards more labeled neurons in the DG of trained mice than in littermate controls (Fig. 13 A-C; Test: 11.62 ± 1.98 Control: 5.84 ± 0.65, p=0.056, Mann-Whitney test, n=5 animals/group). To verify that tdT-labeled cells were recruited during memory recall, animals underwent re-acquisition trials and were sacrificed 90 minutes later for immunostaining for c-Fos. The reacquisition trials identified that the ability of mice to locate the escape box measured by latency to locate the box, the number of errors (Fig. 12). Moreover, the search strategies used to locate the escape box during re-acquisition trials were not different from acquisition day 6 (Fig. 12). Consistently, compared to controls, trained mice showed significantly greater colocalization of tdT with c-*Fos* indicating reactivation of the same cells during memory recall (Fig 11D; Co-Labeling: Test 0.17 ± 0.03 Control 0.03 ± 0.013, p<0.0001 Mann-Whitney test, n=50 sections from 5 mice/group). These results confirm that, unlike task-naïve controls, a significant proportion of DG neurons labeled with tdT in the mice trained on the Barnes maze are activated and reactivated during spatial navigation.

**Figure 13:**
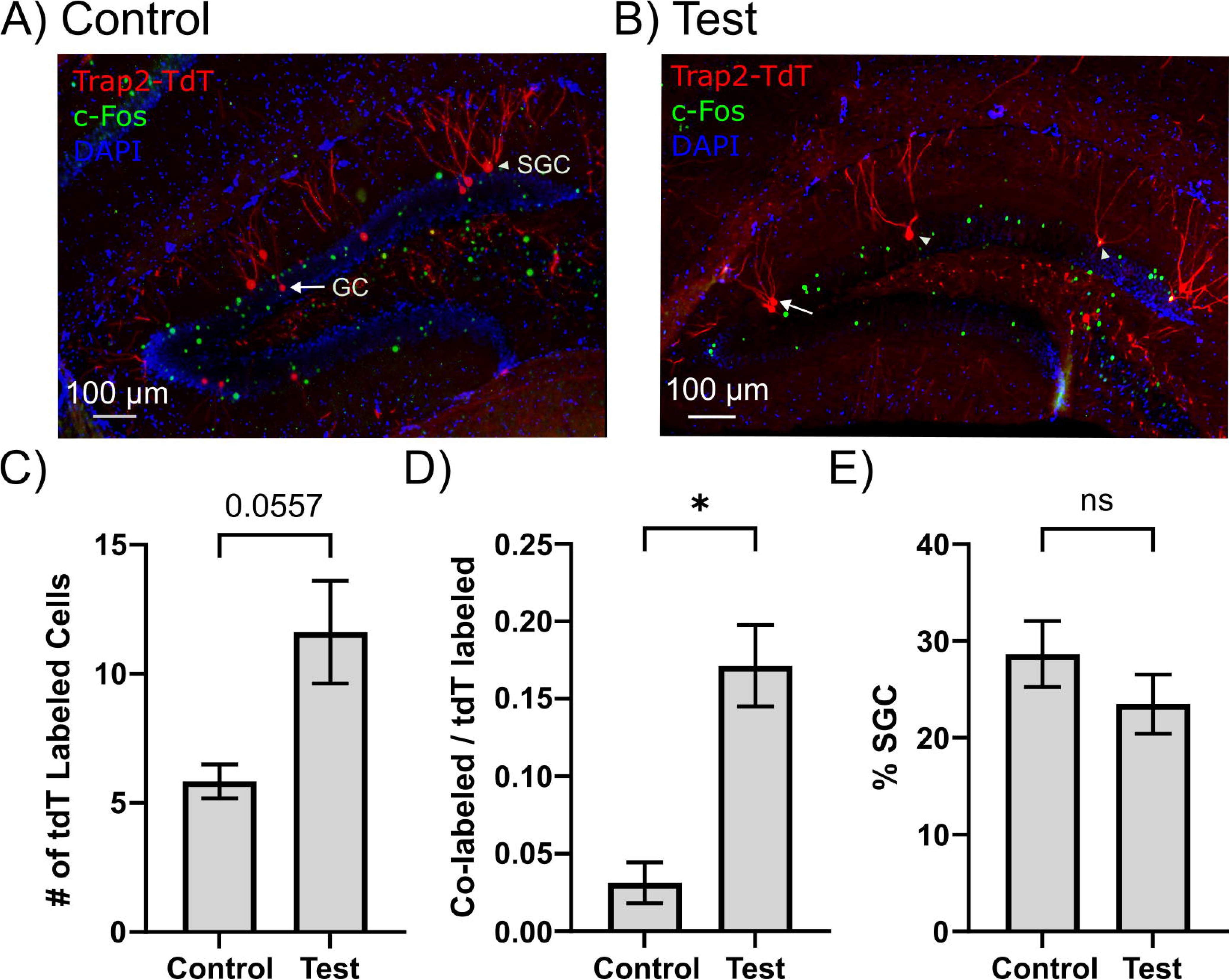
Semilunar granule cells are reliably labeled during c-*Fos*-driven tagging of DG neurons. A-B) Representative epifluorescence images of sections from mice one week after induction of tdT labeling (red) followed by c-*Fos* immunostaining (green) after a reacquisition trial. Arrows highlight morphologically defined GCs, and arrowheads highlight morphologically defined SGCs. Control mice (A) were retained in a home cage and not exposed to behavioral testing. Test mice (B) were treated with tamoxifen on day 6 of acquisition and euthanized after reacquisition for c-Fos labeling. C-D) Quantification of number (C) and proportion (D) of cells labeled with tdT (red) during acquisition that were co-labeled with c-*Fos* (green) following reacquisition (re-aq) E) Proportion of granule cells tagged with tdT that are morphologically classified as SGCs. * indicates p<0.05 by nested t-test. Schematics were generated using BioRender under license.

Having confirmed task-dependent tdT labeling in mice trained in the Barnes maze, we adopted a manual, morphologically based classification of tdT-labeled neurons to quantify the number of labeled GCs and SGCs. SGCs were identified based on the presence of multiple primary dendrites, wide dendritic arbor, and greater soma width compared to height and location in or near the IML (Afrasiabi et al., 2022; Dovek et al., 2025; Dovek et al., 2026). Despite comprising only an estimated 5% of the GC population (Save et al., 2019), approximately 25% of the tdT-labeled cells in task-naïve mice were SGCs, indicating higher basal activity levels in SGCs compared to GCs. Interestingly, the proportion of SGCs labeled as part of active neuronal ensembles was not different between naïve controls and trained littermate mice (Fig. 13E; %SGC: Control 28.65 ± 3.41, Trained: 23.47 ± 3.06; p=0.25, Mann-Whitney test, n=50 sections from 5 mice/ group). These findings suggest that higher basal glutamatergic drive and intrinsic excitability may support disproportionate labeling of SGCs as part of active neuronal ensembles in control mice. Concurrently, the higher proportional SGC labeling is maintained during spatial behavioral tasks, suggesting SGC recruitment during spatial navigation.

## Discussion

The dentate gyrus is a critical node in entorhinal-hippocampal information transfer and plays a role in consolidating inputs from several sensory modalities (Hainmueller and Bartos, 2020; Borzello et al., 2023). Despite its simple trilaminar structure, the DG has extensive molecular and cellular diversity (Dyhrfjeld-Johnsen et al., 2007; Woods et al., 2018; Hainmueller and Bartos, 2020; Yao et al., 2021). Here we demonstrate that SGCs, a morphologically distinct DG projection neuron subtype, receive more frequent spontaneous excitatory synaptic events and have higher overall dendritic spine density than GCs, particularly in the middle dendritic segments that receive MPP input. MPP-evoked EPSC amplitudes in SGCs were greater than in GCs, and MPP activation recruited SGCs more reliably than GCs. While we show that SGCs receive LPP, SuM, and dorsal and ventral commissural inputs, the amplitude of EPSCs evoked by activation of these pathways was not different between cell types. Interestingly, EPSCs in response to LPP activation showed short-term facilitation in GCs and short-term depression in SGCs. Correspondingly, photoactivation of LPP at 10 Hz progressively increased firing probability in GCs but not in SGCs, identifying differential activity-dependent LPP recruitment of GCs and SGCs. Collectively, these data suggest that SGCs and GCs receive differential inputs from ipsilateral input streams with distinct information content, while receiving similar contralateral and SuM inputs. Analysis of passive membrane models of SGCs and GCs revealed that differences in dendritic morphology did not impact dendritic attenuation or synaptic integration between cell types. In addition to previously reported preferential labeling of SGCs as part of active neuronal ensembles during a spatial memory task (Dovek et al., 2025; Dovek et al., 2026), we find greater than expected c-*Fos*-dependent labeling of SGCs even in task-naïve mice. Together, our data demonstrate differences in pathway-specific input strength, short-term plasticity, and recruitment between SGCs and GCs, which position SGCs to be preferentially activated by contextual information from the MEC.

Our demonstration of differences in pathway-and modality-specific recruitment of GCs and SGCs by LEC and MEC is reminiscent of the divergent input streams to superficial and deep CA1 pyramidal cells (Masurkar et al., 2017). Indeed, as with CA1 deep pyramidal cells (Bayer, 1980; Masurkar et al., 2017), SGCs are preferentially generated early in embryonic development, migrate farther radially to the IML (Save et al., 2019), and receive stronger MEC inputs. SGCs also receive stronger associational hilar inputs than GCs, which parallels the difference in the strength of projections from CA2 to deep CA1 pyramidal cells (Kohara et al., 2014). LEC inputs to GCs and SGCs differed not in strength but in short-term dynamics, while commissural projections did not differ between cell types. We also show that, like GCs, SGCs receive SuM inputs and that there are no cell-type-specific differences in the strength of excitatory and inhibitory SuM inputs between GCs and SGCs. Our findings extend the known differences in morphology, intrinsic physiology, developmental timeline, and inhibition between SGCs and GCs (Williams et al., 2007; Larimer and Strowbridge, 2010; Gupta et al., 2012; Save et al., 2019; Gupta et al., 2020; Afrasiabi et al., 2022) and identify diversity in glutamatergic inputs among DG principal cells located along the radial axis. Whether SGCs and GCs differ in molecular and microcircuit features needs further investigation. Beyond mature DG projection neurons, the adult neurogenic niche adds another layer of radial diversity in the dentate (Altman and Das, 1965; Vivar et al., 2012). Adult-born GCs preferentially reside in the inner third of the GCL, near the hilus. Indeed, as in our prior studies (Gupta et al., 2020; Afrasiabi et al., 2022; Dovek et al., 2025; Dovek et al., 2026), we restricted our GC recordings to the middle and outer thirds of the granule cell layer to avoid this immature population. Immature GCs are more excitable and, unlike SGCs, receive stronger LEC synapses than mature GCs (Vivar et al., 2012), indicating an additional layer of cellular complexity in the DG circuit beyond the superficial and deep layer distinction observed in CA1 pyramidal cells. Collectively, dentate projection neuron types that receive different ipsilateral input streams with distinct information content are positioned to enhance the dentate’s computational capacity.

Although the total dendritic length is not different between GCs and SGCs (Afrasiabi et al., 2022), SGCs receive more frequent and larger amplitude spontaneous EPSCs and have a greater spine density than GCs. The higher spine density in SGCs was selective to the proximal and middle dendritic segments, which receive MEC, SuM and commissural/associational inputs. Moreover, the increased spine density in SGCs was predominantly driven by greater thin dendritic spines, previously shown to be associated with learning and synaptic plasticity (Popov et al., 2004; Bourne and Harris, 2007), supporting SGCs’ engagement in memory function. Evaluation of spine types by dendritic segment identified higher thin and mushroom spines in the middle dendrites of SGCs than in GCs, providing a structural framework for preferential SGC recruitment by MPP. Curiously, like mossy cells (Buckmaster et al., 1993) we found that some SGCs have somatic spines and also exhibit spines on putative axons. While our observation of somatic and axonal spines is anecdotal, given the location of SGC somata in the IML, these spines appear ideally positioned to receive commissural, associational and SuM inputs.

Mass activation of the PP elicited EPSCs with comparable amplitude in both GCs and SGCs, as previously reported in rat (Williams et al., 2007). Since MPP and LPP differ in information content and strength of glutamatergic inputs to mature and immature GCs (Schmidt-Hieber et al., 2004; Vivar et al., 2012), we examined these specific input streams using pathway-specific ChR2 labeling. We evaluated glutamatergic currents to isolate synaptic strengths without contamination from feedforward or feedback inhibition or use of synaptic blockers, and evaluated the ability of the inputs to evoke firing in the cell types. To avoid confounding effects arising from variability in AAV-driven reporter expression between animals, and even between sections, we adopted dual SGC-GC recordings at the same depth from the surface. Consistent with the lack of difference in distal dendritic spine density in GCs and SGCs, the LPP-evoked EPSC amplitude and firing probability did not differ between cell types. This contrasts with the stronger LPP-driven EPSCs in adult-born immature GCs compared to mature GCs (Vivar et al., 2012; Woods et al., 2018). Moreover, unlike GCs, SGCs showed short-term depression during a 10 Hz train of LPP-evoked EPSCs, unlike the facilitation seen in LPP synapses in mature and immature GCs (Vivar et al., 2012). As noted above, we excluded immature GCs in this study by recording from GCs in the middle and outer thirds of the GCL with mature dendritic morphology and input resistance less than 400 MΩ (Schmidt-Hieber et al., 2004). The target-specific differences in short-term dynamics of LPP synapses, identified in our study, are similar to reports in CA1 and cortical circuits (Shigemoto et al., 1996; Thomson, 1997; Toth and McBain, 2000). Functionally, given the differences in recruitment during repeated LPP activation, SGCs likely differ from GCs in how they sum and frequency-filter LPP inputs (Quintanilla et al., 2022). In contrast, MPP-driven EPSCs were significantly stronger in SGCs than in GCs, with no difference in multi-pulse depression. SGCs also differ from immature GCs, which show significantly lower MPP-driven EPSC amplitude than mature GCs and short-term facilitation rather than depression (Vivar et al., 2012; Woods et al., 2018). The difference in EPSC amplitude between GCs and SGCs during optical activation of MPP (Fig. 4) contrasts with the lack of cell-type-specific difference in response to mass stimulation of the perforant path (Fig. 2). It is possible that the variability in the proportion of MPP and LPP fibers activated by electrical stimulation in each slice, combined with single-cell recordings and population-averaged analysis, masked the pathway-specific differences in EPSC amplitude that were resolved in paired analysis of data from dual SGC-GC recordings. Taken together, these data suggest that perforant path inputs differentially engage the SGCs, mature GCs and immature GCs for complex information processing. SGCs with strong MPP inputs and short-term depression are ideally suited to be activated by brief contextual information. Mature GCs receive facilitating LEC synapses and depressing MEC synapses (Hsu et al., 2016), which would better enable GCs to integrate sustained LEC activity with contextual information from MEC. Immature GCs receive large-amplitude facilitating LEC synaptic inputs, suggesting that they may strongly respond to object/content information. Because synaptic short-term dynamics can regulate information processing (Tauffer and Kumar, 2021), SGCs are positioned to expand dentate input processing.

Apart from entorhinal inputs, GCs receive commissural and associational glutamatergic projections from long-range mossy cell projections along the septotemporal axis (Buzsàki and Eidelberg, 1981; Frotscher et al., 1991; Hsu et al., 2016; Botterill et al., 2021) and SuM inputs from the hypothalamus (Hashimotodani et al., 2018; Ajibola et al., 2021). Consistent with prior studies (Williams et al., 2007), hilar stimulation of associational fibers elicited larger EPSCs in SGCs than in GCs. Although we took care to eliminate antidromic responses, hilar electrical stimulation could activate both mossy cells and interneurons, which may explain the high response variability. Future studies with transgenic approaches could better isolate associational mossy cell inputs to GCs and SGCs. Regardless, it is worth speculating that associational mossy cell synapses may target somatic spines on SGCs in the IML. Mature and immature GCs receive robust commissural inputs from contralateral mossy cells (Chancey et al., 2014; Abdulmajeed et al., 2022). Similarly, SuM provides excitatory and inhibitory inputs to GCs (Soussi et al., 2010; Hashimotodani et al., 2018; Ajibola et al., 2021). We extend these findings to SGCs, demonstrating that SGCs receive ventral and dorsal hilar commissural synapses as well as SuM inputs that are not different from those in mature GCs.

Neuronal input integration can be influenced by dendritic structure, including branching complexity (Stuart and Spruston, 2015). Consistently, simulations in morphologically constrained passive membrane models of GCs and pyramidal cells have suggested that differences in dendritic branching complexity could underlie the reduced distal dendritic EPSP attenuation in GCs (Jaffe and Carnevale, 1999; Schmidt-Hieber et al., 2007). Both experimental and computational studies have identified prominent EPSP attenuation at GC proximal dendrites (Krueppel et al., 2011). Despite SGCs having lower dendritic complexity than GCs (Gupta et al., 2020; Afrasiabi et al., 2022), our morphologically constrained passive membrane models showed no difference in EPSP dendritic attenuation and summation between GCs and SGCs. However, despite identical channel and leak conductance densities, the same peak synaptic conductance produced larger local dendritic EPSPs in GCs than in SGCs. Differences in membrane surface area may reduce membrane resistance and attenuate EPSP amplitude in SGCs. Consistent with this proposal, the larger somatic compartment size in SGC models contributed to their lower input resistance than in GCs. Similarly, the integration of coincident synaptic inputs within and between branches was less than the numerical sum of individual EPSPs and was not different between cell types. The results in GCs are consistent with the role of active dendritic channels, not included in our passive dendritic model, in mediating linear EPSP summation in GCs (Krueppel et al., 2011). Further experimental evaluation of active dendritic properties and synaptic integration will be needed to determine whether SGCs, support linear summation like GCs or superliner summation like cortical pyramidal cells (Polsky et al., 2004; Krueppel et al., 2011). Regardless, our simulations show that morphological differences do not affect EPSP attenuation or summation in SGCs and GCs.

Unexpectedly, about a quarter of the labeled neurons in both task-naïve controls and mice trained in a spatial memory task were SGCs, well above the estimated 5% SGC population among dentate GCs (Save et al., 2019). The disproportionate labeling of SGCs during a spatial memory task is consistent with prior studies (Erwin et al., 2020; Dovek et al., 2025; Dovek et al., 2026) and with their stronger MEC inputs. While prior studies have not examined SGC labeling in task-naïve controls, we find preferential SGC labeling even in the absence of behavioral exposure. Because ensemble labeling in TRAP2 mice relies on activity-dependent expression of the immediate early gene c-Fos (Yiu et al., 2014; Gouty-Colomer et al., 2016), our finding suggests that SGCs have higher basal activity than GCs, consistent with SGCs having greater spontaneous glutamatergic drive and sustained firing. Another unexpected finding is the similar proportional labeling of SGCs in task-naïve and expert mice, despite using yoked naïve littermates to limit variability in tdT expression in the TRAP2 mice, which indicates task-associated scaling of SGC activity in parallel with GCs during contextual memory formation.

Our demonstration of layer- and modality-specific differences in excitatory inputs to GCs and SGCs suggests that they could enhance the DG’s computational capacity by supporting divergent and parallel input processing and extends the known differences between GCs and SGCs.

## Acknowledgements

We thank Drs. Edward Zagha and Sachiko Haga-Yamanaka for thoughtful discussions. We thank Dr. David Carter at the UCR Microscopy Core and Mr. Erick Contreras for help with imaging. This work is supported by National Institutes of Health (NIH) NINDS R01NS069861, R01NS097750 to V.S., NIH/NINDS F31NS124290 to L.D., AES predoctoral fellowship to C.-W. Y., UCR RISE fellowship to E.G., AES BRIDGES to V.S., and E.G.

## Author contributions

Conceptualization, L.D. and V.S.; Methodology, L.D., C.-W. Y., and V.S.; Investigation, L.D., C.-W. Y., A.T. N, and E.G,; Writing – Original Draft, L.D and V. S.; Writing – Review & Editing, L.D., C.-W. Y., A.T. N, E.G, V.S.; Funding Acquisition, L.D. and V.S.; Supervision, L.D. and V.S.

## Declaration of generative AI and AI-assisted technologies in the writing process

No generative AI and AI-assisted technologies were used in writing this manuscript.

