## Supplementary Figure for "Differential Glutamatergic Inputs to Semilunar Granule Cells and Granule Cells Underscore Dentate Gyrus Projection Neuron Diversity"

### Dovek et al. Supplementary Figures

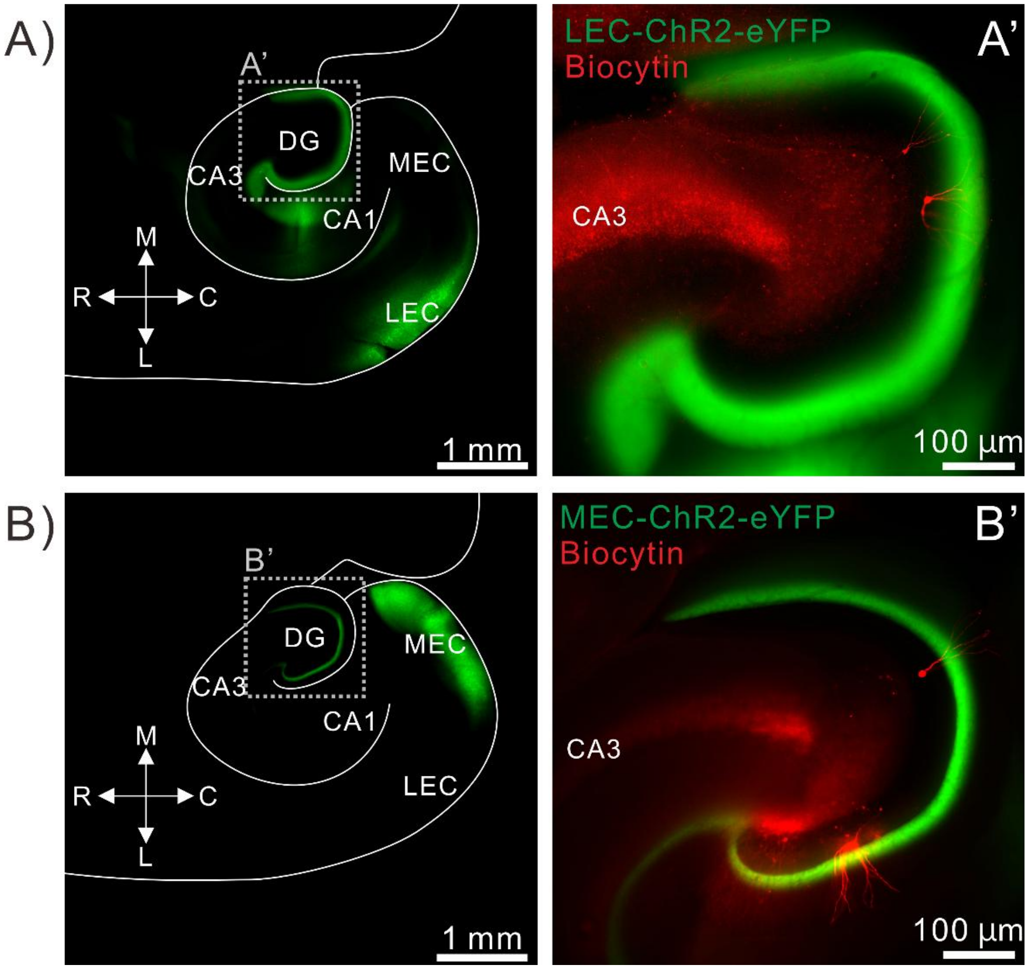


**Supplementary Figure 1: Illustration of LEC and MEC injection sites and pathway labeling**

(A) Representative image of -ChR2-eYFP expression in LEC (green) following AAV injection. (A') A high-magnification image of the DG from the same slice in A illustrates labeling of the lateral perforant path in the outer molecular layer. Biocytin- filled cells and spill in CA3 (red) are shown in A'. (B) Representative image of the ChR2-eYFP expression in MEC following AAV injection. (B') A high-magnification image of the DG from the same slice in B illustrates labeling of the lateral perforant path in the outer molecular layer. Biocytin- filled cells (red) are shown in B'. M: medial, L: lateral, R: rostral, C: caudal, DG: dentate gyrus, LEC: lateral entorhinal co

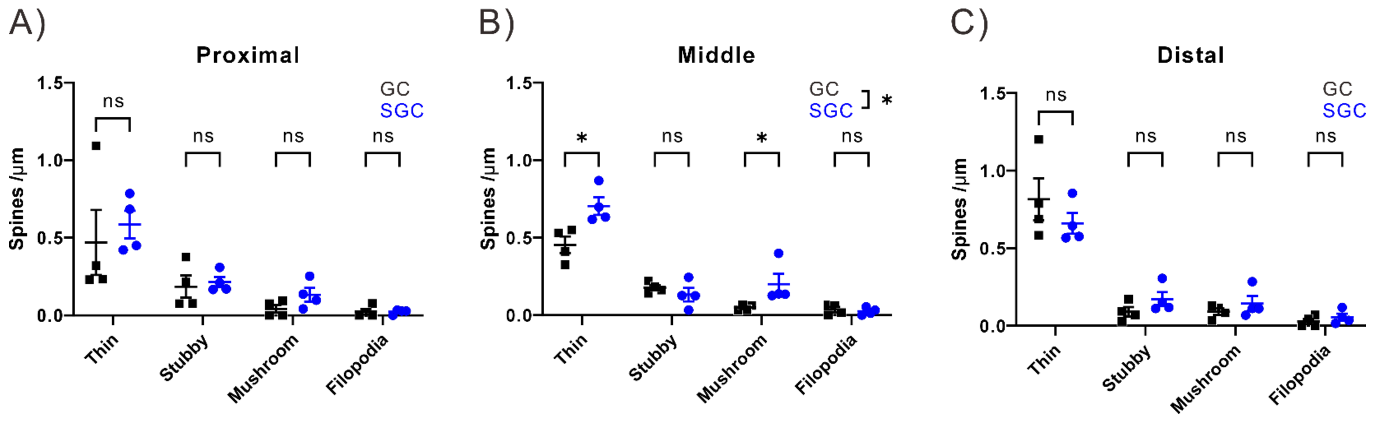


**Supplementary Figure 2: Distribution of spine type by dendritic segment**.

A-C ) Summary plots of densities of thin, stubby, mushroom and filopodial spines in the proximal (A), middle (B) and distal (C) segments in GCs (back, n = 4) and SGCs (blue, n = 4). Cells included here are a subset of cells in Figure 9. * indicates $p<0.05$, by two-way ANOVA with Holm-Šídák's multiple comparisons post hoc test in n=4 cells/group.
