## Supplemental Table for "Differential Glutamatergic Inputs to Semilunar Granule Cells and Granule Cells Underscore Dentate Gyrus Projection Neuron Diversity"

### Dovek et al. Supplementary Tables

SUPPLEMENTARY TABLE 1. Supplementary. Supplementary1. *Summary of spine types by segments*

| | *Cell type* | *P value (cell type)* | *Thin* | *P value* | *Stubby* | *P value* | *Mushroom* | *P value* | *Filopodia* | *P value* |
|---|---|---|---|---|---|---|---|---|---|---|
| *Proximal (IML)* | GC | p=0.4561 | 0.470 ± 0.209 | p=0.8294 | 0.186 ±0.072 | p=0.9666 | 0.042 ± 0.025 | p=0.8528 | 0.024 ± 0.028 | p=0.9899 |
| | SGC | | 0.586 ±0.089 | | 0.215 ± 0.033 | | 0.133 ± 0.045 | | 0.023 ± 0.008 | |
| *Middle (MML)* | GC | *p=0.0235 | 0.455 ± 0.053 | ***p=0.0009 | 0.178 ± 0.017 | p=0.6860 | 0.050 ± 0.009 | *p=0.0436 | 0.036 ± 0.017 | p=0.8453 |
| | SGC | | 0.704 ± 0.057 | | 0.133 ± 0.043 | | 0.200 ± 0.066 | | 0.025 ± 0.012 | |
| *Distal (OML)* | GC | p=0.9743 | 0.816 ± 0.135 | p=0.2899 | 0.090 ± 0.030 | p=0.7246 | 0.091 ± 0.021 | p=0.7890 | 0.028 ± 0.017 | p=0.7890 |
| | SGC | | 0.660 ± 0.067 | | 0.172 ± 0.045 | | 0.144 ± 0.048 | | 0.056 ± 0.022 | |

SUPPLEMENTARY TABLE 2. *Summary of computational simulation data (Figure 11) and statistical analysis by* two-way ANOVA with Holm-Sidak *post hoc* comparison.

| *Stimulation site* | *Cell type* | *Stim site amp. (mV)* | *P value* | *Soma amp. (mV)* | *P value* | $\frac{Soma\ amp.}{Stim\ site\ amp.}$ *(%)* | *P value* |
|---|---|---|---|---|---|---|---|
| *Proximal* | GC | 0.60 ± 0.04 | *p=0.0016 | 0.53 ± 0.03 | *p<0.0001 | 89.01 ± 3.745 | p=0.7329 |
| | SGC | 0.38 ± 0.05 | | 0.30 ± 0.02 | | 83.73 ± 4.16 | |
| *Middle* | GC | 1.18 ± 0.12 | *p=0.0059 | 0.49 ± 0.03 | *p<0.0001 | 46.42 ± 6.11 | p=0.7329 |
| | SGC | 0.67 ± 0.12 | | 0.28 ± 0.02 | | 52.86 ± 6.34 | |
| *Distal* | GC | 2.20 ± 0.14 | *p<0.0001 | 0.47 ± 0.03 | *p<0.0001 | 32.38 ± 5.09 | p=0.7365 |
| | SGC | 0.96 ± 0.12 | | 0.26 ± 0.02 | | 30.43 ± 2.56 | |
